# Directly Measuring Atherogenic Lipoprotein Kinetics in Zebrafish with the Photoconvertible LipoTimer Reporter

**DOI:** 10.1101/2024.05.29.596423

**Authors:** Tabea O.C. Moll, Mackenzie L. Klemek, Steven A. Farber

## Abstract

Lipoprotein kinetics are a crucial factor in understanding lipoprotein metabolism since a prolonged time in circulation can contribute to the atherogenic character of apolipoprotein-B (ApoB)-containing lipoproteins (B-lps). Here, we report a method to directly measure lipoprotein kinetics in live developing animals. We developed a zebrafish geneticly encoded reporter, LipoTimer, in which endogenous ApoBb.1 is fused to the photoconvertible fluorophore Dendra2 which shift its emission profile from green to red upon UV exposure. By quantifying the red population of ApoB-Dendra2 over time, we found that B-lp turnover in wild-type larvae becomes faster as development proceeds. Mutants with impaired B-lp uptake or lipolysis present with increased B-lp levels and half-life. In contrast, mutants with impaired B-lp triglyceride loading display slightly fewer and smaller-B-lps, which have a significantly shorter B-lp half-life. Further, we showed that chronic high-cholesterol feeding is associated with a longer B-lp half-life in wild-type juveniles but does not lead to changes in B-lp half-life in lipolysis deficient *apoC2* mutants. These data support the hypothesis that B-lp lipolysis is suppressed by the flood of intestinal-derived B-lps that follow a high-fat meal.

## Introduction

From insects to vertebrates, lipoproteins shuttle hydrophobic lipids through the hydrophilic environment of the plasma or hemolymph (1, 2). Apolipoprotein B (APOB)-containing lipoproteins (B-lps) are classified by their size, density, and lipid composition. The lipids transported by B-lps, primarily cholesterol esters (CE) and triglycerides (TG), are surrounded by a phospholipid monolayer and apolipoproteins (3, 4). B-lps are sub-classified by their size into chylomicrons (CM), very low-density lipoproteins (VLDL), intermediate-density lipoproteins (IDL), and low-density lipoproteins (LDL), each decorated with one APOB molecule throughout their lifetime (3, 4). While necessary for lipid transport in the bloodstream, elevated levels of B-lps are linked to an increased risk of cardiovascular disease (CVD) through two mechanisms: 1) chronically elevated levels of LDL (5–7) and 2) the duration and intensity of postprandial lipemia resulting from intestinal-derived CM rich in dietary lipids (8–10). Prolonged periods of high plasma LDL are correlated with the accumulation of lipids within endothelial walls (11), initiating plaque formation, immune response, and atherosclerosis (12–14). Both intestinal-derived and liver-derived B-lps are elevated postprandially (15, 16, reviewed by 17), and the degree of elevation can serve as a predictor for future risk of CVD in healthy individuals (18, 19). This postprandial hyperlipemia is hypothesized to be due to various processes, including a reduction of lipoprotein lipase (LPL) activity (20, 21), the saturation of LPL by the influx of CM (16, 22), and the over-secretion of CM (23, 24). Evidently, changes in any of the tightly regulated steps of B-lp synthesis, remodeling, and catabolism can lead to abnormal concentrations of B-lps that influence CVD risk.

Static measures of LDL-cholesterol have been successfully used in clinical settings to evaluate patient CVD risk (25), however, the kinetics of B-lps likely play a major role in CVD disease pathology (26, 27). and due to the difficulties in performing these assays, they are only quantified in a subset of studies. How does B-lps half-life relate to atherogenicity? B-lps that remain in circulation for a prolonged period exhibit more modifications, such as lipid oxidation (28), glycations (29), and de-sialation (30). Specifically, these modifications increase the likelihood of B-lp uptake by endothelial cells (31) and macrophages (32, 33), which promotes plaque formation rather than removal from the circulation (e.g., by the LDL-receptor (LDLR)).

Several techniques have been developed to investigate the factors influencing lipoprotein kinetics in humans. Lipoproteins can be labeled endogenously or exogenously such that are either radioactive or contain stable isotopes (reviewed by (34–36)). Both methods utilize kinetic modeling (37, reviewed by 38), which has advanced significantly over recent years. However, these methods are indirect measures of lipoprotein lifetime, rather arduous to perform, and the modeling assumptions are based on normal populations that could vary unpredictably in disease states. Moreover, isotopic labeling studies have produced difficult to interpret results. In one study, investigators injected radioactive leucine to obtain VLDL-ApoB kinetics while also injecting radioactive palmitate and glycerin to measure VLDL-triglyceride. Using this double-labeling (ApoB and triglyceride), the mean lifetime for plasma VLDL was different depending on radioactive tracer used (39).

A non-invasive method that tags ApoB while circumventing the need for isotopic labeling would be highly advantageous. We exploited the unique attributes of the zebrafish model to develop the first in vivo direct measure of B-lp lifetime, using a photoconvertible protein. The zebrafish is an advantageous model for the study of B-lps. As a vertebrate, the zebrafish shares about 70% of coding genes with humans (40). Many key metabolic genes are conserved, including microsomal triglyceride transfer protein (*mttp*) (41, 42), Ldlr (43), apolipoproteins (*apoBs*, *apoC2*, *apoEs*) (44, 45), and LPL (46). Mttp is required to form and lipidate nascent B-lps in the ER (47). Apolipoproteins have various functions, such as structural proteins (*apoB*, *apoA*) (48), cofactors for LPL to access triglycerides (apoC2) (49), or cofactors for Ldlr to facilitate uptake from the circulation (*apoEs*) (48, 50). Another key gene conserved in zebrafish is the cholesteryl ester transfer protein (cetp) (41, 43–46, 51), the key protein that transfers cholesterol from high-density lipoproteins (HDL) to LDL (52). As a result, the zebrafish has a more human-like lipoprotein profile as compared with some rodents that lack Cetp (53).

The zebrafish as model organism is further auspicious as the zebrafish larva develop externally using the maternally-deposited yolk as the only nutrient source for the first 5 days of life (54). This endogenous food source provides each larva with a consistent nutrient source that is not dependent on feeding, which can be a major source of experimental variability (54). During larval and juvenile development, the larvae are also optically transparent, allowing for the use of in vivo reporters to examine a wide range of physiological processes, which is often not possible, or extremely difficult, in live mammals.

Here, we describe the creation of LipoTimer, a fusion protein of APOB to the photoactivatable fluorescent protein Dendra2 to directly measure B-lp lifetime. This was achieved by inserting the Dendra2 cDNA sequence into the endogenous zebrafish apoBb.1 gene locus (45). Upon UV light exposure, Dendra2 undergoes a cleavage reaction, switching its emission spectrum from green to red wavelengths (55), allowing us to quantify B-lp turnover by the quantification of red-labeled B-lps over time. Using LipoTimer, we found that *pla2g12b* or *mttp* mutations produce small TG-poor B-lps with very short half-lives. These data are consistent with recent work showing the profound resistance to CVD of *Pla2g12^bhlb218/hlb218^* animals(56). In contrast, *ldlr* mutants, similar to what is observed in humans (57), accumulated small B-lps with very long half-lives. We expect that these B-lps kinetic characteristics will profoundly influence CVD risk. Using LipoTimer we also found that juvenile zebrafish fed a high-fat diet present with longer post-prandial B-lp half-life, suggesting that the increased dietary lipid leads to reduced LPL activity and LDLR clearance. Taken together, these data highlight the power of the LipoTimer zebrafish line to better understand the critical role of lipoprotein modifier genes and environmental factors (e.g., diet) on B-lp lifetime.

## Results

### Generation of the ApoB-Dendra2 fusion protein

**I**n general, a static measure from a blood draw is used to determine the plasma B-lp profile and lipid content to infer CVD risk. However, increased levels of B-lps can be the result of changes in B-lp production or catabolism; thus, investigating B-lp kinetics provides key information about B-lp metabolism. Unfortunately, B-lp turnover measures are more difficult to obtain than numbers of circulating B-lps or their associated lipids. Since zebrafish are translucent, we developed an optical method to investigate B-lp turnover in live zebrafish larvae.

A single zebrafish ApoBb.1 molecule (45), like its human ortholog APOB, is incorporated into B-lps in the endoplasmic reticulum (ER) of hepatocytes and enterocytes. Each APOB molecule remains with the B-lp for its entire lifetime (3, 58, 59). Hence, zebrafish B-lp regulation and metabolism can be investigated with a C-terminal ApoBb.1 fusion protein as deployed in the LipoGlo reporter line (60), which is an endogenous fusion of *apoBb.1* to the photon emitting nanoluciferase. B-lp numbers and sizes can be examined and quantified in a single larva after applying the nanoluciferase substrate furimazine to fixed or homogenized animals (60). However, because the substrate is not compatible with fish health nor consistently permeable in living animals, we elected to create a similar reporter using a fluorescent protein, allowing the examination of B-lp in live zebrafish. Using the TALEN-mediated genome engineering method that was used to create the LipoGlo line (60) the photoconvertible fluorophore Dendra2 cDNA sequence was inserted following C-terminus of the *apoBb.1* gene at its endogenous genomic locus. Briefly, one-cell stage embryos were co-injected with mRNA coding a TALEN pair targeting the stop codon of *apoBb.1* and a donor plasmid carrying in-frame sequences for an HA-tag and Dendra2 (Fig. S1A). The injected animals were raised to adulthood, and a germline carrier of the ApoB-Dendra2 fusion was identified after screening the offspring of 13 fish for fluorescence. The fluorescence-positive offspring derived from this founder were raised to create the stable line. Before generating *apoBb.1^Dendra2/Dendra2^* animals, all offspring of the founder were confirmed by PCR (Fig. S1B) and sequencing for evidence of correct integration. These data indicate that the fusion protein was functional (as is the case in LipoGlo animals). Dendra2 B-lps can be visualized under a fluorescent microscope as early as 1 dpf (Fig. 1A), and B-lp numbers can be quantified from the total level of green fluorescence. Green-state fluorescence of ApoB-Dendra2 rises during the first 4 days of development. Thus, B-lp levels increase throughout early development as the yolk lipids are utilized by the yolk syncytial layer (YSL) to synthesize and export B-lps to support the larvae’s rapid development. As development proceeds beyond 4 dpf, green-state fluorescence of ApoB-Dendra2 rapidly decreases. This developmental curve of lipoprotein levels replicates results observed with ApoB-NanoLuc fusion (ref), further indicating this fusion protein is functional. The fluorescence of heterozygous *apoBb.1^Dendra2/+^* larva is, on average, 58 % less bright than that of homozygous *apoBb.1^Dendra2/ Dendra2^* larva (Fig. 1B, Table S1), indicating that the total green fluorescent signal, similar to the LipoGlo luminescence signal, is a direct readout of the B-lp levels and thereby proportional to the levels of ApoBb.1.

**Figure 1:**
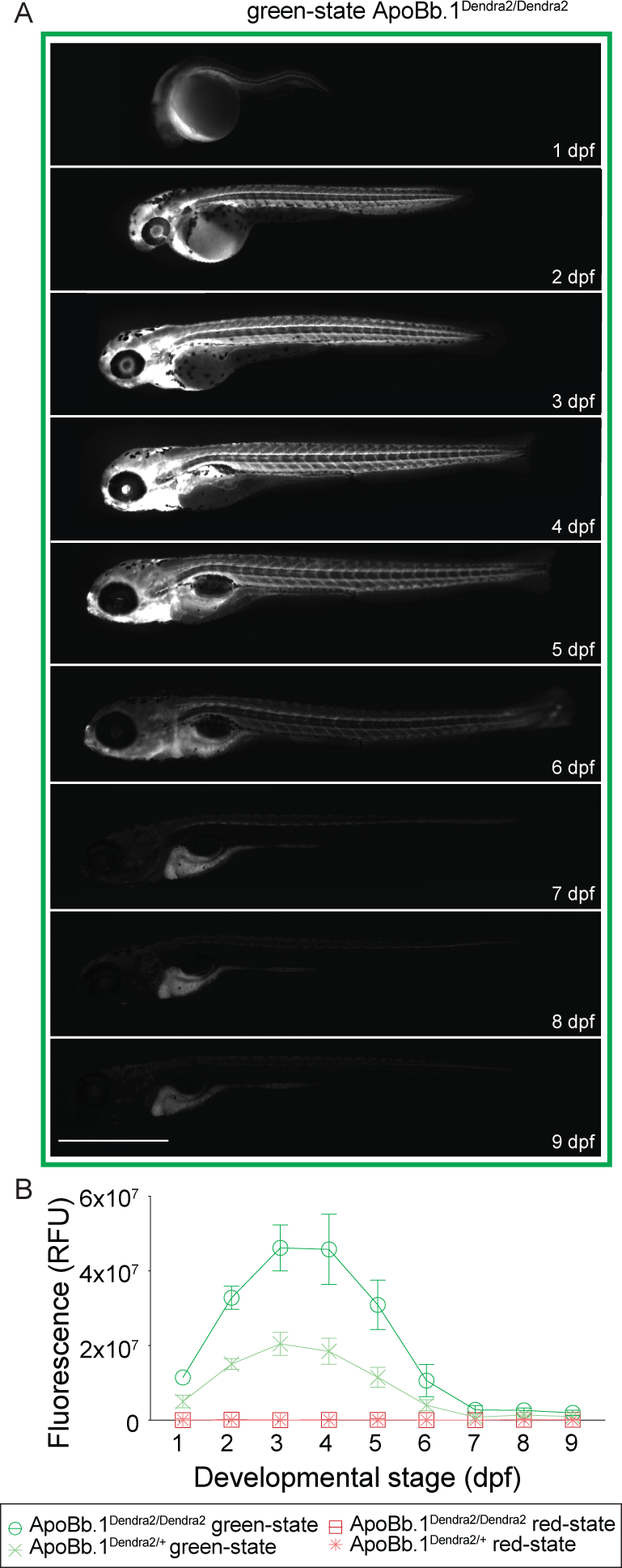
LipoTimer efficiently visualizes B-lps in live animals, and quantification of B-lp levels correlates with yolk utilization. (A) B-lp levels are visualized in LipoTimer larva using green emission filters on a fluorescent microscope. Representative images of the same *apoBb.1Dendra2/Dendra2* larva (1 to 9 dpf) show the localization of the ApoB-Dendra2 signal to the circulatory system and YSL and later the liver. (B) Quantification of green fluorescence from images of *apoBb.1Dendra2/Dendra2* and *apoBb.1Dendra2/+* rise during the first days of development and then fall off to near background levels at 7 dpf. Plotted values reflect the mean of fluorescence after subtraction of the background values of *apoBb.1+/+* animals. *ApoBb.1Dendra2/+* shows half as much fluorescent intensity. Since these animals were not exposed to UV light, no red-state ApoB-Dendra2 is detected. Three independent experiments, n = 15. Scale bar 1 mm.

### Establishing a conversion assay for ApoB-Dendra2 (LipoTimer)

Dendra2 is a photoconvertible, monomeric fluorophore that undergoes conversion when exposed to UV light (55). An irreversible cleavage reaction leads to an emission change from its green-state (Exmax = 490 nm, Emmax = 553 nm) to its red-state (Exmax = 507 nm, Emmax = 573 nm). Thus, B-lps carrying the ApoB-Dendra2 fusion protein are converted to the red-state ApoB-Dendra2 upon UV exposure at a given time point. In contrast, B-lps synthesized after the UV-exposure will only carry green-state ApoB-Dendra2.

To utilize the ApoB-Dendra2 for turnover studies, the photoconversion process must be efficient. Moreover, to measure B-lp lifetime by tracking red-state ApoB-Dendra2, the energy required for photoconversion must neither injure the organism nor alter digestive organ lipid metabolism. To test the impact of UV light intensity on larval health and Dendra2 photoconversion efficiency, 3 dpf *apoBb.1^Dendra2^* larvae were subjected to 0.5 mW/mm^2^ and 2 mW/mm^2^ UV light for up to 2 min in 10 s increments. At each increment, images were taken to evaluate the levels of photoconversion and larval health (Fig. 2A).

**Figure 2:**
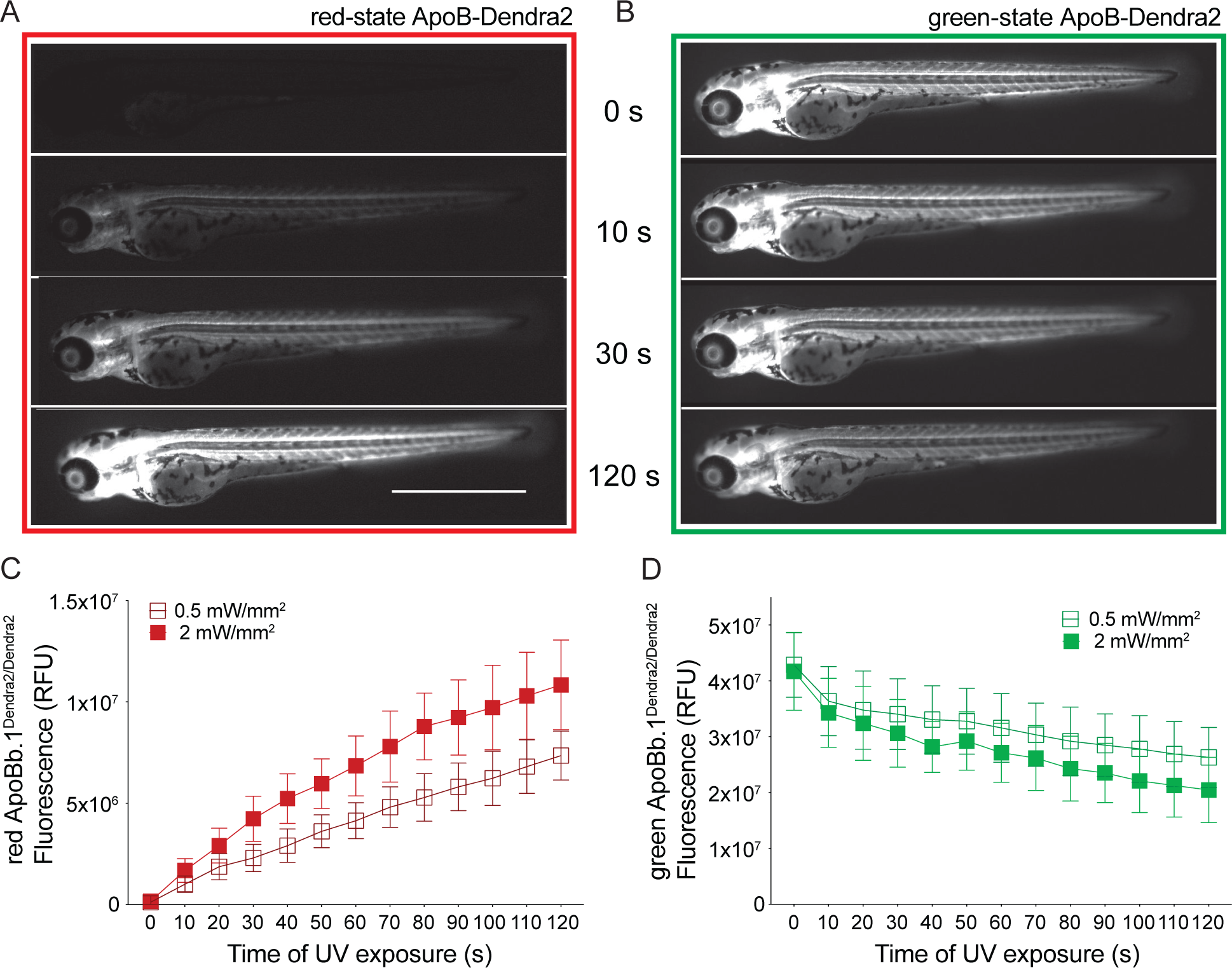
Prolonged UV exposure intensity and/or time increase photoconversion efficiency of ApoB-Dendra2 in larval zebrafish. (A) Photoconversion of ApoB-Dendra2 of 10 s leads to detectable red-state ApoB-Dendra2 levels, which can be increased by prolonged exposure. Representative image of a 3 dpf larva exposed to 2 mW/mm^2^ of UV light. (B) Green-state ApoB-Dendra2 continues to diminish as UV light exposure continues. Representative image of the same larva shown in (A) was obtained by using the green emission filters. (C) The intensity values of red-state ApoB-Dendra2 are plotted as the mean of all imaged larvae, allowing quantification of photoconversion efficiency. Exposure to 0.5 mW/mm^2^ of UV light intensity reduces the photoconversion efficiency. (D) Green-state ApoB-Dendra2 is plotted against time, showing a stronger reduction of green fluorescence in 2 mW/mm^2^ of UV compared to 0.5 mW/mm^2^ of UV. Three independent experiments, each n = 5 of a single clutch, total n = 15. Scale bar 1 mm. When performing the experiment with heterozygous *apoBb.1^Dendra2/+^* animals, photocon-version leads to a 4.1 ± 0.9 fold and 10.74 ± 2.5 fold increase (30 s of 0.5 mW/mm^2^ and 2mW/mm^2^, respectively) of the red-state ApoB-Dendra2 (Fig. S2A), which is 57 – 78 % less than obtained with *apoBb.1^Dendra2/Dendra2^*. All turnover assays were performed using a 30 s UV exposure at 2 mW/mm2 and homozygous *apoBb.1^Dendra2/Dendra2^* animals since these parameters resulted in effective photoconversion while not negatively affecting larval health.

The increase in red-state ApoB-Dendra2 and the concurrent decrease in green-state ApoB-Dendra2 were quantified by measuring the fluorescent signal (Fig. 2 A, B). After 30 s of 0.5 or 2 mW/mm^2^ UV light, an 18.8 ± 5.5 fold and 25.0 ± 6.6 fold increase of red-state ApoB-Dendra2 is measured, respectively, while green-state ApoB-Dendra2 is reduced by 1.3 ± 0.1 fold and 1.4 ± 0.1 fold, respectively. After 2 min of 0.5 or 2 mW/mm^2^ UV light, green-state ApoB-Dendra2 was decreased by 1.6 ± 0.1 fold and 2.0 ± 0.1 -fold, respectively, while red-state ApoB-Dendra2 increased by a striking 60.3 ± 2.6 fold and 64.0 ± 1.8 fold, respectively. However, the health of the larvae was impacted, as evidenced by impaired movement following photoconversion (data not shown) and a drastic increase in auto-fluorescence in the 405 nm channel (Fig. S2C). Auto-fluorescence in the 405 nm channel is indicative of metabolic stress (61) and keratinization of epithelial cells (62). In contrast, these detrimental health effects are not detected using a 30 s UV light pulse of 2mW/mm^2^ (Fig. S2C).

### B-lp turnover accelerates during development

During the first 5 days of development, the larval zebrafish relies on the maternally deposited yolk as the only food source (54). As the embryo develops, the yolk syncytial layer (YSL) forms around the yolk and processes the yolk lipids into B-lps, similar to the intestine and liver (63–65). Towards the end of 5 dpf, the yolk is depleted, and the larval digestive system is developed, so the animal commences feeding. To explore the kinetics of B-lps derived from the yolk, we performed photoconversion of LipoTimer fish at larval ages 2 -5 dpf (Fig. S3A). At each age, homozygous and heterozygous carriers for ApoB-Dendra2 were photoconverted, exposing them to 30 s of 2 mW/mm^2^ UV light. Wild-type siblings, *apoB.1^+/+^*, were included in each experiment as controls for background auto-fluorescence. Following UV photoconversion, red-state ApoB-Dendra2 levels were quantified from individual animals, daily (Fig. 3A and 3B). Example images and fluorescent levels for a larva converted at 4 dpf show a reduction in total red-state ApoB-Dendra2 24 h after photoconversion. Notably, the remaining red-state ApoB-Dendra2 is concentrated at the heart, gills, and the CNS (Fig. 3A). To ensure that turnover measurements derived from this reporter reflect full-length ApoB-Dendra2 turnover and not a breakdown product that might contain only the Dendra2 portion of the fusion protein, we performed Western blots before and after photoconversion and only detected the full-length ApoB-Dendra2 and not free Dendra2 (Fig. S3B and S3C). These data indicated the fluorescent signals from the LipoTimer line reflect the emissions from the full-length fusion protein.

**Figure 3:**
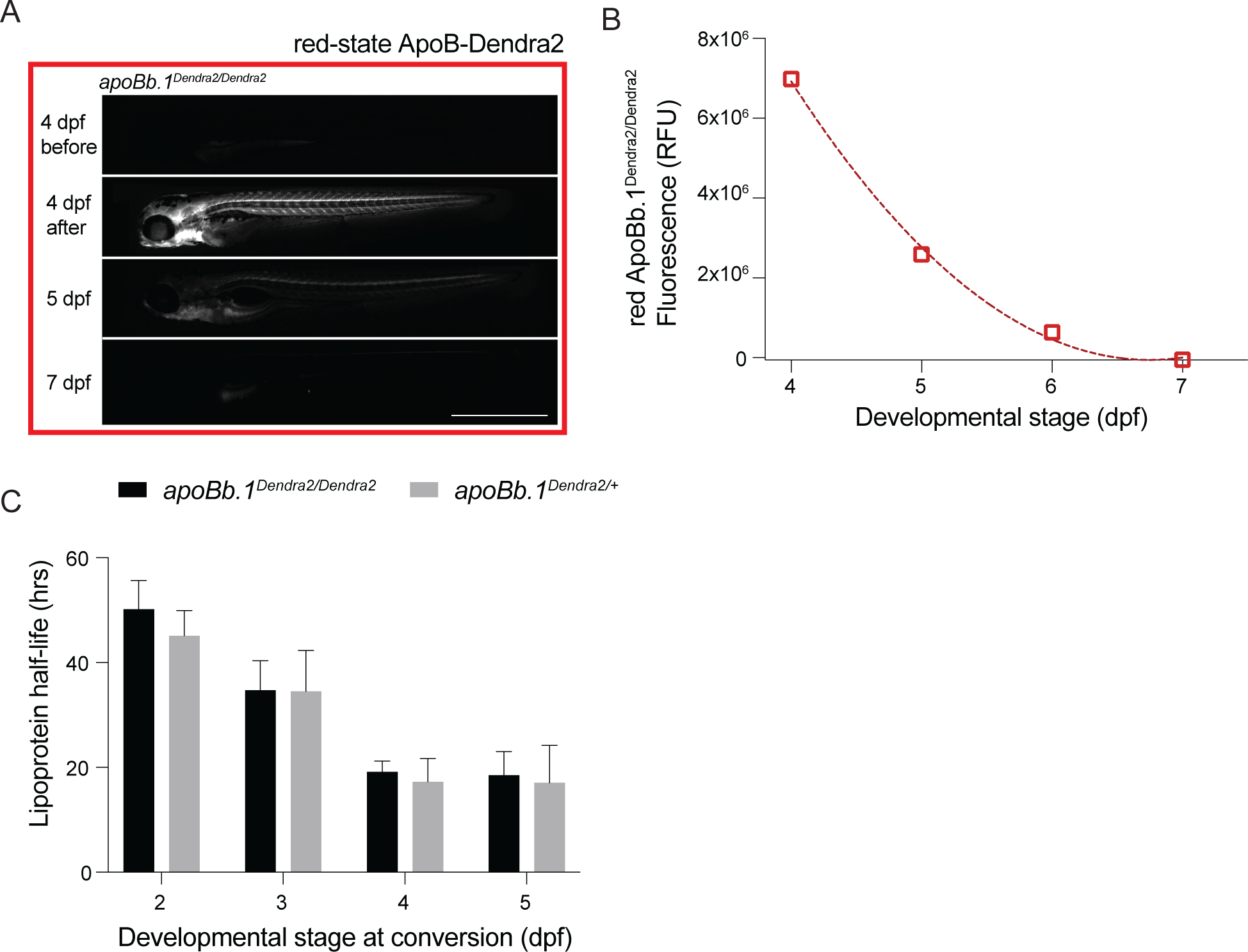
B-lp half-life becomes faster during larval development and is equivalent between heterozygous and homozygous LipoTimer animals. (A) Red-state ApoB-Dendra2 of the larva (4 dpf) increases drastically after photoconversion and diminishes over the course of 72 hrs, as seen in example images of the same animal taken on consecutive days in the red emission channel. (B) The total red-state fluorescent values obtained by quantifying the images with FIJI are plotted and graphed using Prism software. A trend line was fitted to a second-order polynomial equation to calculate the half-life of the red-state ApoB-Dendra2 of each individual larva. (C) Over the course of development, B-lp half-life accelerates from 2 to 5 dpf. Calculation of B-lp half-life in heterozygous and homozygous *apoBb.1^Dendra2^* animals showed no difference between genotypes by Mann-Whitney test. Three independent experiments, n = 7 - 15. Scale bar 1 mm.

We first set out to determine if B-lp turnover was equivalent in heterozygous and homozygous LipoTimer animals during the first 5 days of development. Yolk lipids are packaged into B-lps starting at 2 dpf. When ApoB-Dendra2 is converted at 2, 3, 4, or 5 dpf and red-state ApoB-Dendra2 is subsequently quantified, its half-life declined as development proceeded and no difference between *apoBb.1^Dendra2/Dendra2^* and *apoBb.1^Dendra2/+^* was observed(Table S1). As the half-life measurements between heterozygous and homozygous ApoB-Dendra2 carriers do not show any significant differences, it is concluded that the fusion of Dendra2 to ApoB does not influence the turnover of B-lps.

### Ldlra and apoC2 mutants show longer B-lp half-life

Many of the genes known to profoundly impact plasma lipoproteins are conserved in zebrafish (reviewed by 66, 67-71), such as for the LDLR and APOC2. The LDLR is the main pathway for LDL removal from circulation (72). Patients carrying inactivating mutations of the LDLR gene have significantly higher levels of plasma LDL since uptake is drastically impaired (57). Even patients heterozygous for mutations in LDLR can develop familial hypercholesterolemia and require strict medical and dietary interventions to delay the development of cardiovascular disease (57). APOC2 is a crucial co-factor for LPL, which liberates triglycerides from B-lps (73). Consequently, inactivating APOC2 mutations lead to hypertriglyceridemia (74) and hyperchylomicronemia (75). When mutating the zebrafish ortholog of LDLR, *ldlra^sd52/sd52^*, larvae accumulate lipids in their vasculature, mimicking the human phenotype (43). Similarly, zebrafish mutations in the apolipoprotein C-II (*apoC2*) gene, *apoC2^sd38/sd38^* (44), mimic the human APOC2 plasma lipid phenotype of hypertriglyceridemia (74, 75).

To test the impact of *apoC2* and *ldlra* on total B-lp levels in zebrafish, the *ldlra^sd52^* and *apoC2^sd38^* alleles were crossed into the ApoB-Dendra2 reporter background, and larvae were imaged as they developed (Fig. 4A and 4B). At 5 dpf, increased levels of B-lps were readily detectable in *ldlra^sd52^* and *apoC2^sd38^* mutants when investigating the larvae under the microscope (Fig. 4A and 4B). Quantification of total green-state ApoB-Dendra2 showed significantly increased levels of B-lps in *ldlra^sd52/sd52^* from 4 dpf onwards (Fig. 4 C, Table S1) and *apoC2^sd38/sd38^* larva from 5 dpf onwards (Fig. 4 D, Table S1) compared to their wild-type siblings. From 5 to 7 dpf, we observed an intermediate phenotype of haploinsufficiency as *ldlra^sd52/+^* animals had significantly higher levels of B-lps compared to their wild-type siblings (Fig. 4C, Table S1).

**Figure 4:**
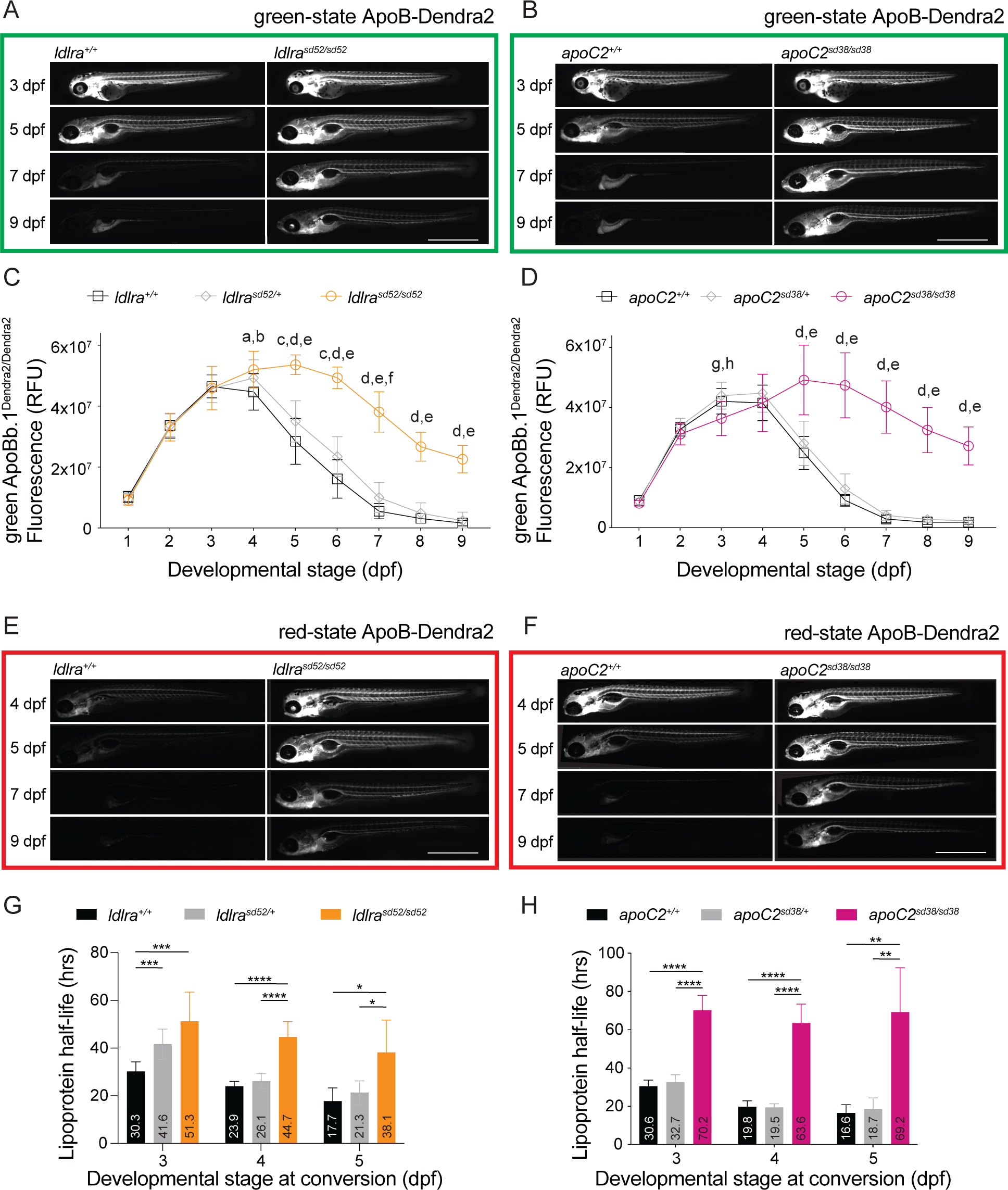
*Ldlra* and *apoC2* mutants have more B-lps starting at 5 dpf and show significantly slower B-lp turnover. Representative images of green-state ApoB-Dendra2 in (A) *ldlra^+/+^* and *ldlra^sd52/sd52^* and (B) *apoC2^+/+^* and *apoC2^sd38/sd38^* larva clearly show the increase in green-state ApoB-Dendra2. Quantification of green-state ApoB-Dendra2 levels in (C) *ldlra* and (D) *apoC2* mutants reveal significantly higher levels of B-lps in the mutants compared to their wild type siblings. 2-Way ANOVA followed by Tukey’s multiple comparisons test; significances as follows a: wild type vs. heterozygous p < 0.01; b: wild type vs. mutant p < 0.01; c: wild type vs. heterozygous p < 0.0001; d: wild type vs. mutant p < 0.0001; e: heterozygous vs. mutant p < 0.0001; f: wild type vs. heterozygous p < 0.05; g: heterozygous vs. mutant p < 0.001; h: wild type vs. mutant p < 0.05. Three independent experiments, n = 6 - 22 (*ldlra*), n = 7 - 31 (*apoC2*). Representative images of (E) *ldlra* and (F) *apoC2* mutants and their wild type siblings that were converted at 4 dpf and imaged on consecutive days. The persistence of the red-state ApoB-Dendra2 is apparent, and when quantified, the B-lp half-life in (G) *ldlra* and (H) *apoC2* mutants is significantly longer. Welch’s ANOVA, Dune’s T3 multiple comparison test; significances as follows *: p < 0.05; ***: p < 0.001; ****: p < 0.0001. Three independent experiments, n = 5 - 28 (*ldlra*), n = 3 - 19 (*apoC2*). Scale bar 1 mm.

To determine the effects of the *ldlra^sd52^* and *apoC2^sd38^* alleles on B-lp half-life, LipoTimer larvae were exposed to UV light and photoconverted from 3 to 5 dpf. These time points were chosen as high numbers of B-lps allow for very efficient photoconversion. Compared to wild-type siblings, increased levels of red-state ApoB-Dendra2 in *ldlra* and *apoC2* mutants were readily detectableas early as 24 hrs post-conversion (Fig. 4E and 4F). Calculation of B-lp half-life showed that B-lps indeed remained in circulation drastically longer in *ldlra^sd52/sd52^* animals at 3 - 5 dpf (Fig. 4G, Table 1, Table S2). Even more striking was the prolonged B-lp half-life in *apoC2^sd38/sd38^* animals at 3 - 5 dpf (Fig. 4H, Table 1, Table S2). In each experiment, mutants were compared to their respective *wild-type* siblings at all ages of conversion (Fig. 4G and 4H). An intermediate phenotype in heterozygous *ldlra* animals was found when measuring the half-life at 3 dpf (p = 0.0002; Fig, 4G, Table 1, Table S2). These data were consistent with our hypothesis that B-lp turnover is drastically decreased when the uptake (*ldlra* mutants) or the lipolysis (*apoC2* mutants) of circulating B-lps is disrupted.

**Table 1:** Whole-body B-lp half-life (in hours)

| Genotype |  | 2 dpf | 3 dpf | 4 dpf | 5 dpf |
| --- | --- | --- | --- | --- | --- |
| *apoBb.1<sup>Dendra2</sup>/Dendra2* | mean ± stdev | 50.2 ± 5.5 | 34.7 ± 5.6 | 19.2 ± 2.0 | 18.5 ± 4.5 |
| *apoBb.1<sup>Dendra2</sup>/+* | mean ± stdev | 45.1 ± 4.8 | 34.5 ± 7.8 | 17.3 ± 4.5 | 17.1 ± 7.1 |
|  | p-value (Mann-Whitney) | 0.0562 | 0.8063 | 0.2312 | 0.3793 |
| *ldlra<sup>+/+</sup>* | mean ± stdev |  | 30.3 ± 3.9 | 24.0 ± 2.0 | 17.8 ± 5.5 |
| *ldlra<sup>sd52</sup>/+* | mean ± stdev |  | 41.6 ± 6.3 | 26.1 ± 3.2 | 21.3 ± 4.9 |
| *ldlra<sup>sd52</sup>/sd52* | mean ± stdev |  | 51.3 ± 12.3 | 44.7 ± 6.5 | 38.1 ± 12.6 |
|  | p-value (Brown Forsythe ANOVA) |  | 0.0002 | 0.0001 | 0.0027 |
| *apoC2<sup>+/+</sup>* | mean ± stdev |  | 30.6 ± 3.2 | 19.8 ± 3.1 | 16.6 ± 4.3 |
| *apoC2<sup>sd38</sup>/+* | mean ± stdev |  | 32.7 ± 3.8 | 19.5 ± 1.9 | 18.7 ± 5.7 |
| *apoC2<sup>sd38</sup>/sd38* | mean ± stdev |  | 70.2 ± 7.8 | 63.6 ± 9.8 | 69.2 ± 22.4 |
|  | p-value (Brown Forsythe ANOVA) |  | 0.0001 | 0.0001 | 0.0001 |
| *mttp<sup>+/+</sup>* | mean ± stdev | 46.1 ± 6.3 | 34.0 ± 5.6 | 23.7 ± 7.3 |  |
| *mttp<sup>c655</sup>/+* | mean ± stdev | 42.7 ± 5.4 | 31.0 ± 3.6 | 21.3 ± 3.0 |  |
| *mttp<sup>c655</sup>/c655* | mean ± stdev | 32.8 ± 7.6 | 24.5 ± 3.4 | 17.6 ± 2.8 |  |
|  | p-value (Brown Forsythe ANOVA) | 0.0080 | 0.0001 | 0.0959 |  |
| *pla2g12b<sup>+/+</sup>* | mean ± stdev | 53.0 ± 7.7 | 38.6 ± 5.9 | 26.9 ± 6.2 |  |
| *pla2g12b<sup>sa659</sup>/+* | mean ± stdev | 54.2 ± 9.9 | 41.4 ± 5.7 | 30.4 ± 9.1 |  |
| *pla2g12b<sup>sa659</sup>/sa659* | mean ± stdev | 41.3 ± 14.6 | 27.6 ± 6.0 | 18.9 ± 3.1 |  |
|  | p-value (Brown Forsythe ANOVA) | 0.0466 | 0.0001 | 0.0007 |  |

### Mttp and pla2g12b mutants present with shorter B-lp half-life

Mttp is an essential protein for B-lp assembly (47, 76). As APOB is translated into the ER lumen, the MTP complex, consisting of MTTP and protein disulfide isomerase (77, 78), binds APOB and co-translationally begins the formation of a B-lp by loading phospholipids and triglycerides onto APOB with the help of Phospholipase A2 group XII B (PLA2G12B) (47, 76). In the absence of MTP, APOB is degraded, and B-lps cannot be formed (47, 56, 76), leading to severe steatosis due to lipid retention (79, 80). Thus, only severe cases of hypercholesterolemia are treated with MTP inhibitors (81, 82).

The zebrafish *mttp^c655^* allele exhibits poor triglyceride transfer and near wild-type levels of phospholipid transfer activity (83). Consequently, B-lps produced by *mttp^c655^* animals are triglyceride-poor and smaller in size. This is in contrast to a more severe zebrafish allele of MTP *mttp^stl^* that produces extremely few lipoproteins (83). Moreover, because some lipid transfer is retained in *mttp^c655^* animals, they do not exhibit the severe liver and intestine lipid retention phenotypes observed in *mttp^stl^* or when animals are treated with the MTP inhibitor lomitipide (83).

Similar to *mttp^c65^* mutants, a small B-lps particle size phenotype was observed in zebrafish with a mutation in *pla2g12b*. Recent work described Pla2g12b as a novel co-factor for MTP (56). Contrary to its name, Pla2g12b does not possess any phospholipase activity but is involved in the expansion of nascent B-lps in the ER (56, 84). The *pla2g12b^sa659^* allele shows an accumulation of lumenal ER lipid droplets and inhibited B-lps synthesis (56). Most intriguingly, mouse mutants for *Pla2g12b* are completely resistant to CVD using the PCSK9 overexpression model of atherosclerosis (56). We hypothesized that these two mutations would result in faster B-lp turnover.

Total B-lp levels of the *mttp^c655^* and *pla2g12b^sa659^* mutants were measured in the LipoTimer reporter background, and larvae were imaged from 1-9 dpf. Images of the animals showed an overall reduced ApoB-Dendra2 signal starting at 5 dpf, and the remaining ApoB-Dendra2 signal was localized to the CNS, heart, and gills (Fig. 5A and 5B). It was notable that *pla2g12b^sa659/ sa659^* mutants presented with an increased ApoB-Dendra2 signal in the YSL, the site of B-lp synthesis. Whole-body B-lp quantification showed that at 4 dpf, *mttp^c655/c655^* mutants carried fewer B-lps than their wild-type siblings, which became more prominent by 6 dpf (Fig. 5C, Table S1). Similarly, *pla2g12b^sa659/sa659^* mutants had less total B-lps at 4 - 6 dpf (Fig. 5D, Table S1). After 6 dpf, no differences between *mttp^c655/c655^* and *pla2g12b^sa659/sa659^* mutants and their wild-type or heterozygous siblings were detectable (Fig. 5C and 5D, Table S1), consistent with B-lp levels previously obtained from LipoGlo assays (83, 85).

**Figure 5:**
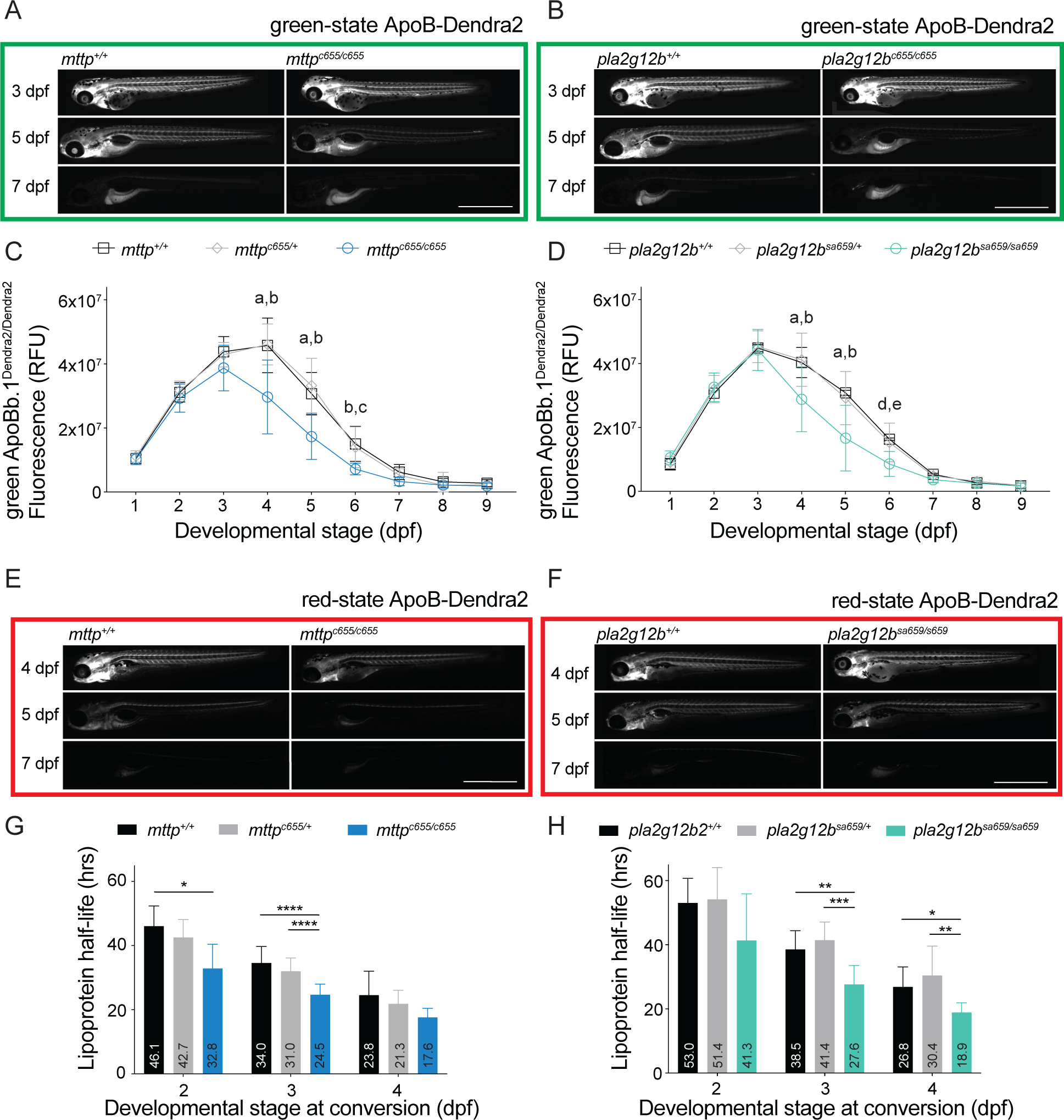
*Mttp* and *pla2g12b* mutants have less B-lps at 4 and 5 dpf and show significantly faster B-lp turnover. (A) A mild reduction in overall green-state ApoB-Dendra2 can be seen at 5 dpf in *mttp^c655/c655^* mutants compared to their *mttp^+/+^* siblings. (B) At 4 dpf, the green-state ApoB-Dendra2 signal is more intense in the YSL of *pla2g12b^sa659/sa659^* mutants than their *pla2g12b^+/+^* siblings. Also, *pla2g12b^sa659/sa659^* mutants have less overall green-state ApoB-Dendra2 in their circulation at 5 dpf compared to *pla2g12b^+/+^* larva. Quantification of green-state ApoB-Dendra2 levels in (C) *mttp* and (D) *pla2g12b* mutants reveal significantly lower levels of B-lps in the mutants compared to their wild type siblings at 4 to 6 dpf. 2-Way ANOVA followed by Tukey’s multiple comparisons test; significances as follows a: wild type vs. mutant p < 0.0001; b: heterozygous vs. mutant p < 0.0001; c: wild type vs. mutant p < 0.01; d: wild type vs. mutant p < 0.001; e: heterozygous vs. mutant p < 0.001. Three independent experiments, n = 11 - 29 (*mttp*), n = 15 - 22 (*pla2g12b*). (E) Representative images of 4 dpf *mttp^+/+^* and *mttp^c655/c655^* larva showing a distinct reduction of red-state ApoB-Dendra2 in *mttp^c655/c665^* with barely discernible fluorescent signal at 5 dpf. (F) Red-state ApoB-Dendra2 in *pla2g12b^sa659/sa659^* is mildly reduced in mutants compared to their wild type siblings when converted at 4 dpf and imaged on consecutive days. The persistence of the red-state ApoB-Dendra2 in the YSL can be made out. When calculating the B-lp half-life of animals converted at 2, 3, or 4 dpf, (G) *mttp* mutants showed a shorter B-lp half-life at all days while (H) *pla2g12b* mutants only had significantly shorter B-lp half-life at 3 and 4 dpf of conversion. Welch’s ANOVA, Dune’s T3 multiple comparison test; significances as follows *: p < 0.05; **: p < 0.01; ***: p < 0.001; ****: p < 0.0001. Three independent experiments, n = 6 - 24 (*mttp*), n = 6 - 18 (*pla2g12b*). Scale bar 1 mm.

B-lp turnover was measured at 2, 3, and 4 dpf in the *mttp^c655/c655^* and *pla2g12b^sa659/sa659^* mutants since the total levels of B-lps were so low after 4 dpf in the mutant animals that conversion efficiency was low and difficult to track. A quick reduction in red-state ApoB-Dendra2 was noticed in *mttp^c655^* mutants (Fig. 5E). *Pla2g12b^sa659^* mutants did not have a drastic decrease in circulating red-state ApoB-Dendra2 but showed retention of the signal in the YSL as seen before in the green-state ApoB-Dendra2 (Fig. 5F). The calculated half-life for *mttp^c655^* mutants showed a significant reduction at 2 and 3 dpf between mutants and their heterozygous and wild-type sibling controls which was mild at 2 dpf and was more pronounced at 3 dpf (Fig. 5G, Table 1, Table S2). In contrast, *pla2g12b^sa659^* mutants had a significantly shorter B-lp half-life at 3 and 4 dpf, but not at 2 dpf. (Fig. 5H, Table 1, Table S2). Accumulation of both green and red state ApoB-Dendra2 in the YSL, suggests that a portion of newly synthesized and converted B-lps remain trapped in this tissue after synthesis. In a recent publication, the *pla2g12b^sa659^* mutation was shown to prevent efficient B-lp secretion, leading to an accumulation of B-lps in the ER (56). Based on this data, we hypothesized that there could be two populations of APoB-Dendra2 with different half-lives: those that fail to leave the YSL (long half-life) and those secreted into the circulation (short half-life). To better measure the circulating portion, we developed an alternative analysis method.

### Circulating lipoproteins in mttp and pla2g12b mutants confirm shorter B-lp half-life

For a specific readout of B-lp half-life in circulation, we developed a targeted analysis to quantify B-lp-associated fluorescence from a specific area of larvae. A 45-pixel wide circle above the anus that excludes all B-lp-producing tissues (e.g., YSL, liver, and intestine) was chosen as the region of interest (ROI) for fluorescent signal quantification to exclude any signal that may result from B-lps not yet secreted and to minimize variation between larvae (Fig. 6).

**Figure 6:**
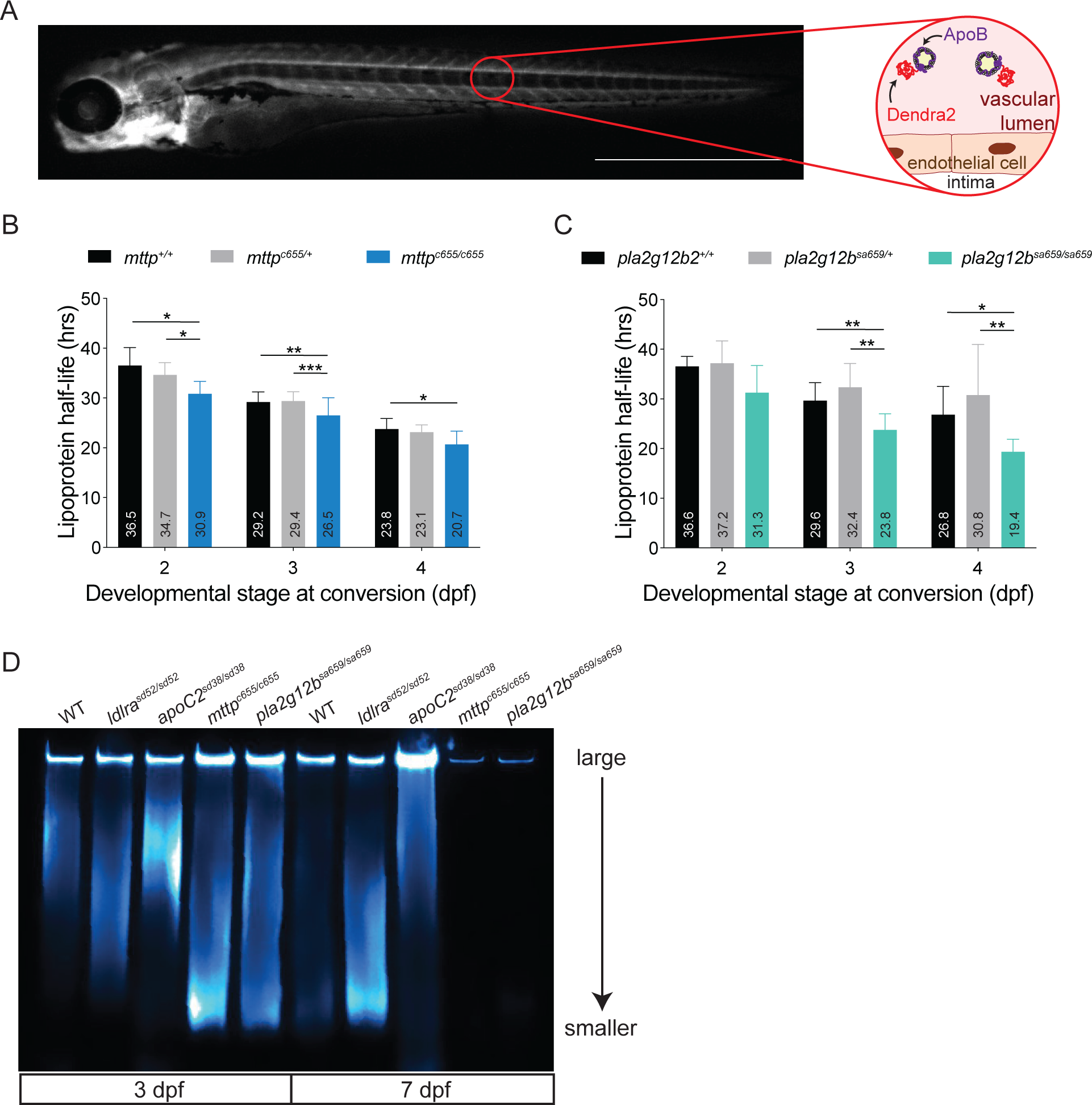
Circulating B-lps are turned over faster in *mttp* and *pla2g12b* mutants, which only produce small B-lps. (A) Representative image of a 4 dpf larva imaged for red-state ApoB-Dendra2. The red circle indicates the area where fluorescence was measured to calculate the B-lp turnover of circulating B-lps, as explained by the circular insert. (B) Circulating B-lps show a shorter half-life in *mttp^c655/c655^* mutants compared to their wild-type siblings when converted at 2, 3, or 4 dpf. (C) B-lp half-life was significantly shorter in *pla2g12b^sa659/sa659^* compared to their wild-type siblings when converted at 3 and 4 dpf but not when converted at 2 dpf. Welch’s ANOVA, Dune’s T3 multiple comparison test; significances as follows *: p < 0.05; **: p < 0.01; ***: p < 0.001. Three independent experiments from Figure 5 were reanalyzed: n = 7 - 30 (*mttp*), n = 8 - 21 (*pla2g12b*). Scale bar 1 mm. (D) Representative LipoGlo-Electrophoresis at 3 and 7 dpf of wild type, *ldlra*, *apoC2*, *mttp*, and *pla2g12b* mutant larvae. At 3 dpf, *ldlra* mutants appear similar to wild types, while *apoC2* mutants carry only large B-lps, and *mttp* and *pla2g12b* mutants only have small B-lps. At 7 dpf, wild-type animals have only a few remaining small B-lps, and *ldlra* mutants have many small B-lps left. *ApoC2* mutants still exhibit only large B-lps, and *mttp* and *pla2g12b* mutants have such low numbers of B-lps that they cannot be detected on the assay. n = 2 from one clutch.

We found that *mttp^c655^* mutants have a shorter circulating B-lp half-life compared to their wild-type siblings at all developmental ages investigated (Fig. 6B, Table 2, Table S3), similar to whole-body B-lp turnover rates discussed above (Fig. 5). While *pla2g12b^sa659^* mutants also mimicked the whole-body B-lp half-life, with no significant change in B-lp turnover at 2 dpf, but shorter half-life of circulating B-lps at 3 and 4 dpf (Fig. 6C, Table 2, Table S3) compared to whole-body B-lp half-life.

**Table 2:**
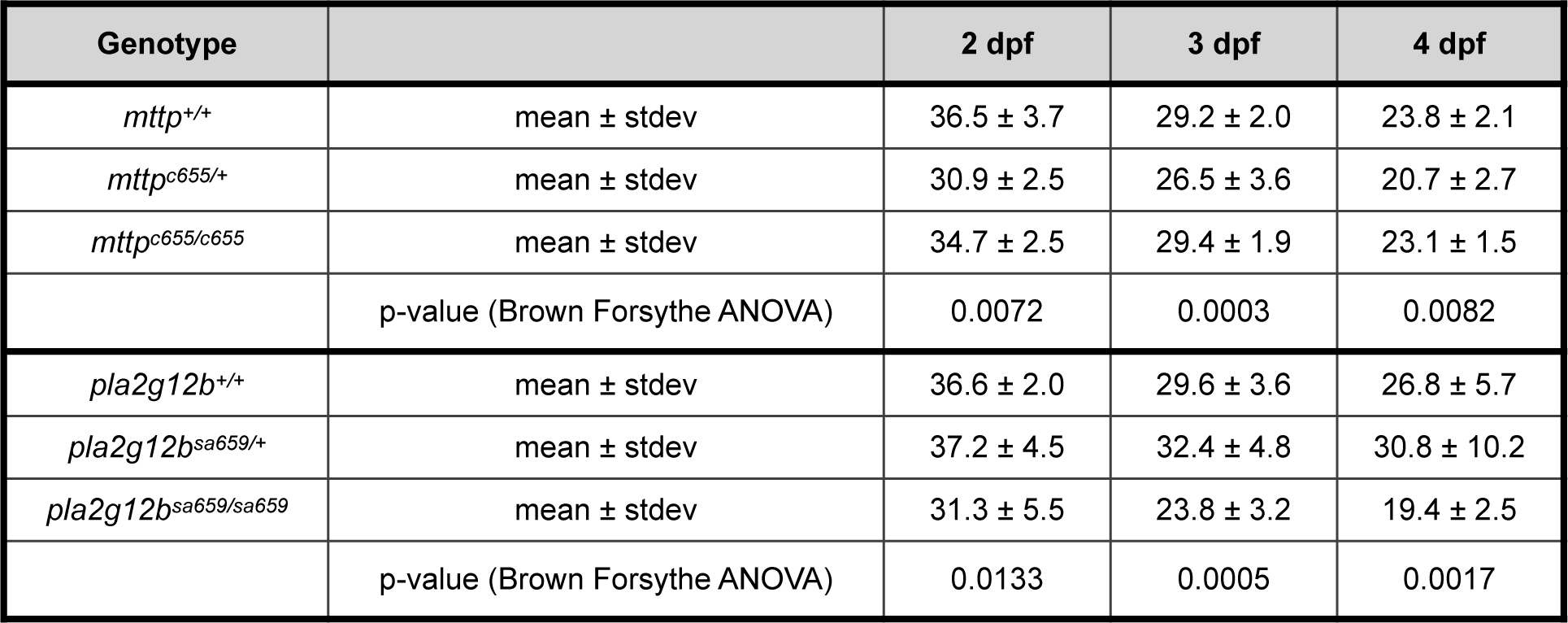
Circulating B-lp half-life (in hours)

| Genotype |  | 2 dpf | 3 dpf | 4 dpf |
| --- | --- | --- | --- | --- |
| *mttp*<sup>+/+</sup> | mean ± stdev | 36.5 ± 3.7 | 29.2 ± 2.0 | 23.8 ± 2.1 |
| *mttp*<sup>c655/+</sup> | mean ± stdev | 30.9 ± 2.5 | 26.5 ± 3.6 | 20.7 ± 2.7 |
| *mttp*<sup>c655/c655</sup> | mean ± stdev | 34.7 ± 2.5 | 29.4 ± 1.9 | 23.1 ± 1.5 |
|  | p-value (Brown Forsythe ANOVA) | 0.0072 | 0.0003 | 0.0082 |
| *pla2g12b*<sup>+/+</sup> | mean ± stdev | 36.6 ± 2.0 | 29.6 ± 3.6 | 26.8 ± 5.7 |
| *pla2g12b*<sup>sa659/+</sup> | mean ± stdev | 37.2 ± 4.5 | 32.4 ± 4.8 | 30.8 ± 10.2 |
| *pla2g12b*<sup>sa659/sa659</sup> | mean ± stdev | 31.3 ± 5.5 | 23.8 ± 3.2 | 19.4 ± 2.5 |
|  | p-value (Brown Forsythe ANOVA) | 0.0133 | 0.0005 | 0.0017 |

We hypothesized that both size and half-life impact the atherogenic character of B-lps. To test this, we used the LipoGlo reporter to determine the size profile of B-lps in larval zebrafish(60). All sizes of B-lps are seen in wild-type animals at 3 dpf. By 7 dpf, the yolk is absorbed, and no external food was provided, leading to a few small B-lps remaining in wild-type larvae (Fig. 6D). When investigating *ldlra^sd52^* mutants, we found a normal B-lp size distribution at 3 dpf compared to the wild-type larva (Fig. 6). However, a drastic accumulation of small B-lps can be seen at 7 dpf in *ldlra^sd52^* mutants, consistent human mutations of Ldlr that lead to the accumulation of LDL in circulation (57) (Fig. 6 D). In contrast, *apoC2^sd38^* mutants present with a large B-lp-phenotype at 3 dpf that is still persistent at 7 dpf (Fig. 6 D). Intriguing, however, are the B-lp profiles seen in *mttp^c655^* and *pla2g12b^sa659^* mutants. Both present with a small B-lp phenotype at 3 dpf, similar to the 7 dpf *ldlra* mutant profile. At 7 dpf, *mttp^c655^* and *pla2g12b^sa659^* mutants have such low levels of B-lps that no B-lp profile can be visualized (Fig. 6D).

### B-lp half-life is longer in juvenile zebrafish fed a high-cholesterol diet

Numerous studies have shown that the consumption of a high-fat meal increases not only CMs but also liver-derived B-lps (15, 16, 22, 86). Newly synthesized CMs, which comprise most of the total B-lps following a meal (23, 24), monopolize LPL (16, 22) and outcompete the already circulating B-lps, likely extending their lifetime in circulation. We wanted to test this hypothesis by adapting the larval B-lp turnover assay to juvenile zebrafish to perform a direct measure of B-lp lifetime in response to a high-cholesterol diet (8 % cholesterol-supplemented standard food). Since the standard GEMMA diet consists of 14 % lipids with a basal level of 0.6 % cholesterol (87), spirulina was chosen as the low-fat diet (7 % lipids and minimal cholesterol (88)) and has been studied as a zebrafish diet (89–91). We defined the high-cholesterol diet, as standard GEMMA food supplemented with 8% cholesterol, using a recently developed method that leads to hypercholesterolemia in larval zebrafish (87). Juvenile zebrafish were housed in individual enclosures, called playpens so that they could be repeatedly measured throughout the experiment (92). Juveniles were converted at 15 dpf, at which age they can consume larger food but are still easily imaged. Zebrafish fed the standard diet have variable B-lp levels (Fig. S4A). To generate a consistent and measurable population of B-lps (Fig. S4A), ApoB-Dendra2 fish were fed a Western diet for 1 day before photoconversion (Fig S4, Fig. 7A). One hour after the morning feed at 15 dpf, the animals were removed from the tank and randomly chosen for the two feeding conditions (Fig. 7A). Animals were separated into four tanks, two dedicated for low-fat diet and two for high-cholesterol diet. Each tank held 12 playpens, 9 with 1 *apoBb.1^Dendra2/Dendra2^* each, 3 with 1 control *apoBb.1^+/+^* animal each. All tanks were fed their respective diet three times daily after the photoconversion procedure. Before feeding separate diets, all animals had similar B-lp levels (Fig 7B). Feeding the high-cholesterol diet led to a 1.48-fold increase in total B-lp numbers at 16 dpf (p = 0.0008), which became more prominent by 20 dpf (4.91-fold increase, p < 0.0001; Fig. 7B). The increased cholesterol led to overall larger juveniles at the end of the experiment (low-fat diet 117.5 ± 97.7 µm length vs. high-cholesterol diet 324.3 ± 196.8 µm length; p < 0.0001; Fig. 7C). Animals fed a low-fat diet had a significantly lower B-lp half-life compared to high-cholesterol diet fed animals, consistent over 3 experiments (low-fat diet 44.0 ± 16.0 h vs. high-cholesterol diet 54.3 ± 18.63 h, p < 0.0227) (Fig. 7D). When comparing the individual rates of B-lp half-life to the growth of the fish, no correlation was detected (Fig. S5), indicating that the increased growth is not the driver of the change in B-lp half-life.

**Figure 7:**
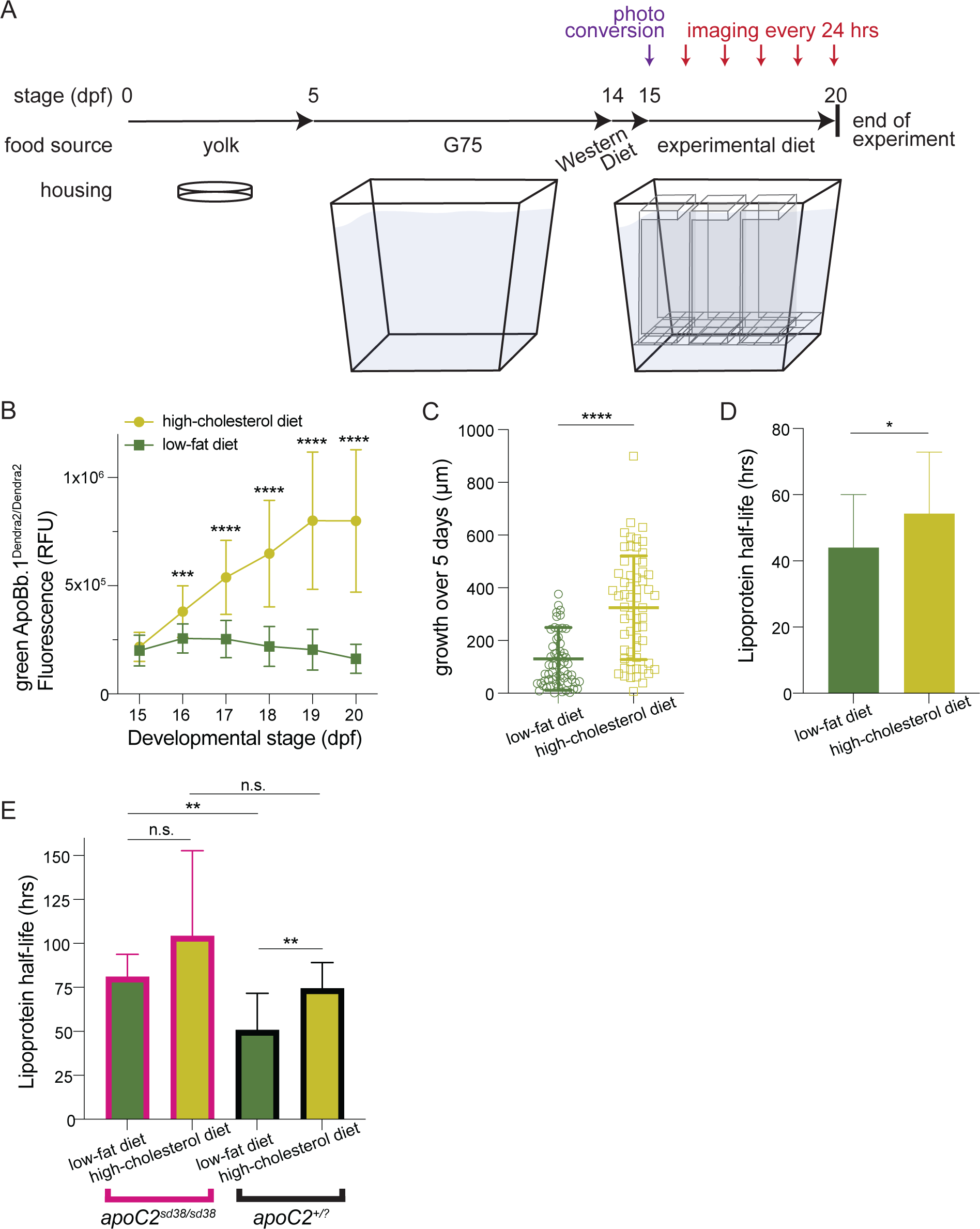
Feeding a high-cholesterol diet increases circulating B-lp numbers and juvenile growth and prolongs B-lp half-life in wild-type animals, but not in *apoC2* mutants. (A) Schematic of feeding experiments: larvae are raised in dishes until they commence feeding in a 10 L tank on the recirculating system. After one day of feeding a Western diet, 15 dpf old juveniles are photo-converted and moved into playpens in 10 L tanks, receiving their respective experimental diet. (B) Feeding a high-cholesterol diet leads to significantly more circulating B-lps than the low-fat diet. ***: p = 0.0008, ****: p < 0.0001. 2way ANOVA followed by Šídák’s multiple comparisons test. (C) Juveniles fed a high-cholesterol diet exhibit more growth from day 1 to day 5 of the experimental feeding. ****: p < 0.0001. Mann-Whitney test. (D) Feeding a high cholesterol diet from 15 - 20 dpf leads a longer B-lp half-life in wild-type juveniles compared to siblings fed a low-fat diet. * p = 0.0227. Mann Whitney test. Three independent experiments, n(total) = 46-50 (E) *apoC2* mutants fed the low-fat diet exhibit increased B-lp half-life compared to their wild-type and heterozygous siblings. ** p = 0.0082. However, there is no change in B-lp half-life in *apoC2* mutants in response to the high-cholesterol diet, while wildtype and heterozygous siblings have a longer B-lp half-life on the high-cholesterol diet compared to the low-fat diet. ** p = 0.0026. Mann Whitney test. Three independent experiments, n(total) = 3-16.

To examine if LPL activity was the driver for the increased B-lp half-life, *apoC2^sd38^* mutants and their siblings were fed a high-cholesterol diet. Since the larval experiments did not indicate a heterozygous phenotype, *apoC2* wild-type and heterozygous animals were pooled into the *apoC2* control group in the juvenile analysis, annotated ad *apoC2^+/?^*. When fed the low-fat diet, *apoC2^sd38^* mutants had a slower turnover than their wild-type and heterozygous siblings (*apoC2^sd38^* mutants 81.1 ± 12.6 h vs. *apoC2^+/?^* controls 50.8 ± 20.8 h, p = 0.0082) (Fig. 7E). However, feeding the high-cholesterol diet led to similar B-lp half-life in all *apoC2* genotypes (*apoC2^sd38^* mutants 104.4 ± 48.3 h vs. *apoC2^+/?^* controls 74.42 ± 14.7 h, p = 0.3648) (Fig. 7E). Intriguingly, *apoC2* mutants had no difference in B-lp half-life on the low-fat and high-cholesterol diet (low-fat diet 81.1 ± 12.6 h vs. high-cholesterol diet 104.4 ± 48.3 h, p = 0.5714) (Fig. 7E). These data are consitent that the increased B-lp half-life on a high-cholesterol diet results of LPL oversaturation.

## Discussion

In 1950, the association between lipoproteins and CVD prompted a surge in research dedicated to lipoproteins (93, 94). Soon after, lipoprotein kinetic studies utilizing radioactive isotopes to investigate the influence of lipoprotein kinetics on hypercholesterolemia were developed (95). Since then, clinical practice has evolved to predominantly focus on static measures of lipoprotein and cholesterol levels to evaluate a patient’s risk for CVD (25). These parameters are relatively straightforward to obtain, whereas studying the lifetime of a B-lp in circulation is more challenging. Current methods in humans and animal models label B-lps with tracers and then determine their lifetime by analyzing blood draws and applying models to calculate B-lp half-life (35, 37, 96). The LipoTimer zebrafish reporter is the first direct method to measure B-lp turnover via imaging of live zebrafish.

The endogenous fusion on the C-terminus of *apoBb.1* aims to label only lipidated B-lps, as ApoB undergoes ER-associated decay when it is not co-translationally lipidated (47, 76). Considering that ApoB must be lipidated to form a lipoprotein and we observe a single ApoB-Dendra2 fusion protein expressed in animals by Western blot (Fig. S3 B, C), we expect that the fusion is stable and functional. Further, homozygous carriers of the transgene did not exhibit a darkened yolk, a phenotype observed in *apoBb.1^wz25/wz25^* mutant animals (97), nor did they exhibit altered growth, behavior, or fecundity. By using minimal exposure to UV light to photoconvert B-lps to red-state, we observed no significant detrimental health effects on the animals. We observed that heterozygous LipoTimer animals present with 58 % less signal of green B-lps compared to homozygous carriers suggesting a slight preference for the wild-type ApoBb.1 during the formation of B-lps. We hypothesize a preference for the *wild-type* allele in heterozygous animals as the addition of a bulky tag may partially reduce. Further, we noticed that photoconversion is 57 - 78 % less effective in heterozygous *apoBb.1^Dendra2/+^* animals compared to homozygous *apoBb.1^Dendra2/Dendra2^*, in line with a preference for the wild-type ApoB protein., Regardless, the conversion process demonstrates comparable effectiveness in both genotypes. The B-lp distribution is visually equal between control animals and animals that were exposed to UV light, and the B-lp lifetime is the same in heterozygous and homozygous ApoB-Dendra2 animals. Both support that B-lp characteristics are not altered by the Dendra2 fusion to ApoB and the photoconversion process.

B-lp half-life in wild-type larvae significantly changes during larval development. B-lps produced at 2 dpf remain in circulation for much longer than B-lps produced at 3, 4, or 5 dpf (Fig. 3). The shortened half-life later in development is likely attributable to the interplay of many factors: During later developmental stages, the rapidly developing animal requires more nutrition to support the increased body size, simultaneously the yolk begins to rapidly diminish such that fewer B-lps are produced, and during these first 5 days exogenous food is not an option because the digestive organs have not finished developing. Zebrafish mutants for *ldlra* and *apoC2* both have increased numbers of B-lps, mimicking the human disease phenotypes of hypercholesterolemia and hypertriglyceridemia (57, 75). While *ldlra* mutants accumulate small B-lps and *apoC2* mutants experience an abundance of large B-lps, both mutations lead to a significantly slower B-lp turnover. In contrast, the *mttp^c655^* and *pla2g12b^sa659^* mutants show a significantly faster B-lp turnover. This is of particular interest as these mutants consistently produce small, triglyceride-poor B-lps that are attributed to have a negative impact on vascular health (98, 99). Contrary to this expectation, these mutants exhibit no increase in vascular endothelial lipid deposit (83). However, we cannot yet draw any conclusions on the effect of these mutations on atherosclerosis in zebrafish, as the zebrafish community has yet to establish a reproducible, reliable model for intravascular lipid deposition associated with cardiovascular disease (56, 100). Instead, the mouse model was utilized to evaluate the effect of *Pla2g12b* atherosclerosis (56). In the context of LDLR-reducing PCSK9 overexpression, together with a chronic Western diet, *Pla2g12b* mutant mice do not develop any lipid depositions in the aortic arch, which are readily detectable in the sibling controls. It appears that the shorter B-lp half-life in the *Pla2g12b* mutants may indeed be a key regulator of atherosclerosis in this context (Chapter 4, Fig 5).

While it is not yet possible to investigate the half-life of one specific B-lp subgroup, the ApoB-reporter allows the measurement of B-lp kinetics in the entire animal, the circulation, or any region of interest visible in a fluorescent image. In the *mttp* and *pla2g12b* mutants, the B-lp half-life in the entire animals was slightly longer compared to the B-lp half-life of the circulating population. Likely, the whole-body B-lp half-life is longer due to including B-lps in the quantification that are still being synthesized and exported.

Intriguingly, we also detect green-state and red-state ApoB-Dendra2 in the central nervous system (CNS). A similar CNS localization of B-lps was observed using the LipoGlo reporter (60). The Reissner fiber, a structure in the cerebral spinal fluid, is populated by the protein Scospondin (101). Mutants for Scoscpondin develop severe scoliosis (101), and the loss of its LDL-binding domain could be speculated as a causative factor. Images of red-state ApoB-Dendra2 show a prolonged signal in the CNS, allowing us to hypothesize that the LDL-binding domain of Scospondins might trap B-lps instead of taking them up. The importance of the LDL-binding domain of Scospondins is the subject of ongoing future studies.

Additionally, we adapted the LipoTimer assay to examine the impact of dietary changes on circulating B-lps turnover rates. We chose two diets for this examination: 1) a low-fat diet consisting of spirulina, a food source of very low lipid content, and 2) a high-cholesterol diet, prepared according to (87), with 8 % cholesterol added to the conventional diet. Our high-cholesterol diet had recently been described to increase levels of B-lps and, eventually, lead to steatosis (87). Indeed, when fish were fed the high-cholesterol diet, they showed increased levels of circulating B-lps compared to animals fed the low-fat diet and grew to a larger overall size. No correlation between zebrafish size and B-lp turnover could be detected, suggesting B-lp half-life is independent of growth at this stage. Since *apoC2* mutant juveniles on both diets present with similar rates of B-lp turnover, our data support the hypothesis that postprandial hyperlipidemia is a result of LPL saturation (16, 20–22).

With the LipoTimer assay, it is possible to investigate the changes in B-lp half-life in response to various diets, and the effects of short- and long-term diet-induced hyperlipemia. However, some animals showed an increase in circulating red-labeled B-lps at the 24 h post-conversion time point. Previous research suggests that the primary release of postprandial CMs occurs within 45 – 60 min (102). Consequently, the juvenile conversion experiment was performed 60 – 90 min after the morning feed. However, the animals were allowed to freely feed for 60 min and the ingestion of food could have been closer to the conversion time point for each individual (example see Fig. S5C). Furthermore, the time frame of primary CM release was obtained from humans and could be drastically different in zebrafish. Hence, we hypothesize that newly synthesized CMs are converted and released into the bloodstream, leading to the observed increase in circulating B-lps at the 24 h time point. To account for the increase and still allow the calculation of B-lp half-life in these animals, the curve fit was adapted to a third-order polynomial. In animals displaying the 24 h increase, the first derivative of the third-order polynomial was calculated and used to determine half-life. Likely due to the overall slower uptake of B-lps in *apoC2* mutants, the post-conversion red B-lp increase was more prone in *apoC2* mutants (87 % of *apoC2^sd38/sd38^* vs. 46 % of *apoC2^+/?^* on the low-fat diet and 90 % of *apoC2^sd38/sd38^* vs. 70 % of *apoC2^+/?^* on the high-cholesterol diet).

In addition to genetic and diet perturbations, it will be interesting to apply this new method to study the effects of drugs on B-lp kinetics. An interesting first experiment would include PCSK9 inhibitors, which decrease LDLR degradation after endocytosis and thus increase B-lp clearance (103). Kinetic studies with human participants revealed that PCSK9 inhibitors showed both reduced production of B-lps and increased LDL turnover (104).

Overall, the new LipoTimer reporter allows for the first direct *in vivo* examination of B-lp kinetics. We found that B-lp half-life is drastically longer in zebrafish mutants for *ldlra* and *apoC2*. In contrast, mutants for *mttp* and *pla2g12b,* which only produce small, triglyceride-poor B-lps, present with increased B-lp turnover. These data indicate the importance of B-lp kinetics as a contributor to lipoprotein atherogenicity. In a final experiment, we showed that high levels of post-prandial lipids lead to a slower B-lp turnover, supporting the hypothesis that LPL saturation is a key driver of post-prandial hyperlipidemia.

## Acknowledgments

We gratefully acknowledge Julia Baer, Mackenzie Klemek, and Hannah Kozan for fish husbandry. We thank Dr. McKenna Feltes and Dr. Daniel Kelpsch for helpful editing assistance. We also thank Dr. Mahmud Siddiqi for support with microscopy and the development of the photoconversion process. This work was supported by The National Institutes of Health grants F31HL149174 (T.O.C.M.) R01DK093399 (S.A.F.), R01GM63904 (S.A.F.) and the Carnegie Institution for Science endowment (S.A.F.). The content is solely the authors’ responsibility and does not necessarily represent the official views of the National Institutes of Health (NIH).

## Author Contributions

T.O.C.M. performed experiments, analyzed data, prepared figures, and wrote the manuscript. M.L.K. (under the supervision of T.O.C.M) performed conversion experiments. S.A.F. designed and supervised experiments and contributed to the writing and revision of the manuscript.

## Materials and Methods

### Zebrafish husbandry

Adult zebrafish were fed once daily with ∼3.5% body weight Gemma Micro 500 (Skretting USA). Natural spawning of group crosses with two females and two males provided embryos, which were kept in embryo medium at 28 °C on a 14:10 light: dark cycle until 6 days post fertilization (dpf) unless noted otherwise. Larvae were then moved to the fish facility (14:10 light: dark cycle) and fed with GEMMA Micro 75 (Skretting) three times a day until 14 dpf unless noted otherwise. Then, food was changed to GEMMA Micro 150 twice daily and Artemia once daily from 15 - 42 dpf.

### Genome editing

In-frame integration of Dendra2 into the C-terminus of apoBb.1 was achieved using TALENs as previously described and validated (56) (Addgene stock 128695 and 128696). TALENs were *in vitro* transcribed using the T3 Message Machine Kit (Thermo Fisher Scientific, AM1348). A donor plasmid was obtained from Genewiz, designed to include homology arms for the TALENs with a left homology arm of 537 bp in the coding sequence of apoBb.1 and a right homology arm of 754 bp in the 5’ UTR of apoBb.1. Between those sequences, an in-frame HA-tag, followed by a 4-glycine linker, and the Dendra2 coding sequence was inserted to serve as the template for DNA integration (Addgene stock TBD). One-cell stage embryos obtained by natural crosses of Casper/Casper adults were co-injected with an injection mix including 150 ng/ µL of TALEN mRNA and 12 ng/µL of donor plasmid (Fig. S1). Injected embryos were raised to adulthood, and progeny were crossed to AB and screened for fluorescence of Dendra2.

### Quantification of ApoB-Dendra2 levels using microscopy

Larvae were mounted in 3% methylcellulose/embryo media and imaged using the AxioZoom V16 microscope, with a Zeiss Plan NeoFluar Z 1x/0.25 FWD 56 mm objective, AxioCam MRm camera, and Zen 2.5 software at 25 x magnification. The exposure time for both channels was set to 50 ms, and acquisition ROI was set to 1388 x 232 pixels. Juveniles were imaged at 16x magnification, with an exposure time of 150 ms, and acquisition ROI of 1388 x 600 pixels.

After the initial imaging, larvae were moved into EM in a 24-well plate for housing during the experiment to allow for the tracking of each individual larva over time.The image’s total fluorescence intensity was obtained using FIJI and its measurement of “Integrated Intensity”. In each experiment, 3 - 4 *apoBb.1^+/+^* larvae were imaged and averaged as background signals. Using the attached Excel template 1, all values were transcribed, and the intensity of each larvae was calculated.

### Photoconversion of apoBb.1^Dendra2^ larva

Larval and juvenile zebrafish were immobilized in 3% methylcellulose and placed under a 385 nm high-power LED light (Thor Labs, SOLIS-385C). The LED Driver (Thor Labs, DC2200) tuned the LED light intensity and duration. Fold-change increase was calculated relative to the mean of the initial timepoints. To photoconvert Dendra2 from green to red, larval and juvenile zebrafish were placed under the LED light for 30 s at 50 % intensity.

### Measuring and calculating B-lp lifetime

Experiments started at 10:00/10:30 AM and ran until 11:00/11:30 AM to ensure consistent age-staging. Before photo conversion, each fish was imaged using the above-mentioned settings to obtain the green and red fluorescence baseline. Fish were imaged immediately after photo conversion and placed in individual wells of a 24-well plate. Fish were imaged every 24 hrs until larvae reached 9 dpf or red levels of Dendra2 were no longer detectable. Using FIJI, the total levels of lipoproteins were calculated by measuring “Integrated Intensity”. All values were inserted into the attached Excel template 2. *ApoBb.1^+/+^* sibling controls were included in every experiment; average intensity for these animals was used for background subtraction.

In larval experiments, the red intensity values for each fish were fitted to second-order polynomial equations in Prism.

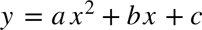

The B0 (c), B1 (b), and B2 (a) values and R squared were used to calculate the half-life in Excel. Animals with an R^2^ < 0.9 were excluded from the calculation of half-life, which excluded, on average, in 3 % of the larval experiments.

In juvenile experiments, the background of the last imaging day was removed equally from all images. Each fish’s red fluorescent values were then fitted to a third-order polynomial equation in Prism.

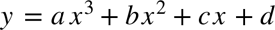

With Prism delivering B0 (d), B1 (c), B2 (b), and B3 (a). To calculate the half-life of the juveniles, the maximum of the polynomial fit was determined using the first derivative of the third-order polynomial and the half-life was then attributed half of the curve maximum. To solve for the equation, the internet calculator WolframAlpha was used. R-squared values were obtained from Prism as well, and animals with an R2 < 0.9 were excluded. Additionally, animals with no real solution were exluded, this led to overall X % of fish excluded in wild-type experiments and X % of fish excluded in the *apoC2* experiments.

When measuring the B-lp half-life of circulating lipoproteins, a circular ROI of 45 px was placed on the fin above the larva’s rectum. The Integrated Intensity of the ROI was measured, and values were then processed the same way as for data from whole-body calculations.

Scripts were written for the facilitation of data analysis in FIJI and are provided as supplemental files.

### DNA extraction and genotyping

The HotSHOT DNA extraction protocol was adapted to obtain genomic DNA (105). Adult fin clips were heated to 95 °C for 20 min in 50 µL and single larva in 20 µL of 50mM NaOH. After cooling to room temperature, 10% volume of 1 M Tris pH 8.0 was added.

Integration confirmation of the Dendra2 coding sequence and general genotyping for *apoBb.1^Dendra2^* was performed with primers 1, 2, and 3 (T_a_ = 57 °C, extension time = 20 s), with the WT band at 113 bp and the successful fusion at 369 bp. For sequencing the entire integration site, primer 4 on apoBb.1 but not on the donor plasmid was paired with primer 2 (T_a_ = 57 °C, extension time = 30 s).

For conversion assays, natural spawning of group crosses was performed with adults *gene-of-interest^+/-^*; *apoBb.1^Dendra2/+^*. At 1 dpf, embryos were sorted by fluorescent intensity for *apoBb.1^Dendra2/Dendra2^* and *apoBb.1^+/+^*. After finishing the conversion assay, larvae were genotyped with specific primers (0.5 µM concentration unless noted otherwise) for each *gene-of-interest* (Supplementary Table 1). The *ldlra^sd52^* locus was genotyped with primers 5 and 6 (T_a_ = 64 °C, extension time = 30 s), with a WT band of 128 bp and a mutant band of 118 bp. The *apoC2^sd38^* deletion was genotyped with primers 7 and 8 (T_a_ = 65.5 °C, extension time = 20 s), leading to WT 147 bp and mutant 137 bp. The *mttp^c655^* mutation used primers 9 and 10 (T_a_ = 50 °C, extension time = 30 s) as described in (83). The PCR product was then digested using the BsrI restriction enzyme (3 units of BsrI (New England Bio-Labs, R0527) in New England BioLabs Buffer 3.1 (B7203)) as the mutation introduces a cut site, leading to a WT band of 137 bp and mutant band sizes of 76 bp and 61 bp, and heterozygous with all three bands post digestion (65 °C, 3.5 h). Primers 11 and 12 (T_a_ = 57 °C, extension time = 30 s) were used for genotyping the *pla2g12b^sa659^* mutations described in (60). The PCR product was digested (37 °C, 3 hrs) with Bts1-v2 (3 units of Bts1-v2 (New England Bio-Labs, RR0667S) in New England BioLabs Buffer rCutSmart Buffer (B6004S)). The mutation introduces a cut site, leading to a WT band of 150 bp and mutant band sizes of 111 bp, and heterozygous with both bands.

### LipoGlo-Electrophoresis

See (60) for detailed LipoGlo methods and reagents (Promega Corp., N1110; (106)). Larvae carried one copy of the LipoGlo reporter (*apoBb.1^Nluc/+^*). For homogenization, larvae were dispensed into 96-well plates (USA Scientific, #1402-9589) in a total volume of 100 µL of B-lp stabilization buffer (40 mM EGTA, pH 8.0, 20% sucrose + cOmplete Mini, EDTA-free protease inhibitor (Sigma, 11836170001)) and sonicated in a microplate-horn sonicator (Qsonica Q700 sonicator with a Misonix CL-334 microplate horn assembly). Homogenate was stored at -20 °C until use. Native-PAGE was prepared according to (60) and imaged using the Odyssey Fc (LI-COR Biosciences) gel imaging system. The Di-I LDL standard migration assigned areas for Zero Mobility (ZM), VLDL, IDL, and LDL.

### Western blot

For protein extraction, 10 larvae were pooled per condition. *ApoBb.1^Dendra2/Dendra2^* larvae were snap-frozen before and after conversion at 3 dpf. At 5 dpf, *ApoBb.1^Dendra2/Dendra2^* larvae converted 48 hrs prior, and non-converted siblings were snap-frozen, along with *ApoBb.1^+/+^*. To transfer and thaw larvae, 100 µL of 1x RIPA buffer (10X RIPA (Millipore Sigma, 20-188) diluted and 1 cOmplete, Mini Protease Inhibitor Cocktail (Millipore Sigma, 11836153001) was added to the larvae. Larvae were homogenized using a pellet pestle and incubated at 4 °C for 15 min while shaking. Samples were centrifuged for 5 min at 12,000 x g at 4 °C, and the supernatant was mixed with equal volume 2 x Laemlli Buffer (Bio-Rad, 1610737) and heated to 95 °C for 5 min. For Dendra2-only control, the Dendra2 protein was obtained from OriGene Technologies, Inc (TP790047), and 5 µL of Dendra2 was added to 95 µL of 1 x RIPA and 100 µL 2x Laemlli buffer. Precision Plus Protein All Blue Prestained Protein Standards (Bio-Rad, 1610373) was used as a molecular weight marker. A precast 4 - 20 % gradient gel (Bio-Rad, 4561093) was loaded with 25 µL of each sample and 10 µL of Dendra2 protein and run for 30 min at 70 V and 60 min at 90 V. Transfer buffer was prepared (125 mL 4x Running Buffer, 825 mL deionized water, 50 mL methanol, 500 µL 10% SDS, 560 µL ß-mercaptoethanol) and pre-chilled before overnight protein transfer at 30 V constant onto a PVDF. Coomassie stain was performed to confirm protein transfer. After blocking the blot in 5 % milk for 1 hr at room temperature, primary mouse monoclonal antibody binding Dendra2 (Thermo Fisher, TA180094, 1:400 dilution) was incubated overnight at 4 °C. The membrane was washed three times in HRP wash (6.05g Tris, 8.75g NaCl in 1 L deionized water, pH 8), then incubated for 1 hr with an goat anti-mouse HRP-antibody (EMD Millipore, #12-349, diluted 1:10,000). After three more washes with HRP wash, imaging solution (Fisher Scientific, PI34095) was added, and the membrane was imaged in the chemi channel for 2 min using the LiCor Fc(Odyssey) Imager.

### Diet preparation

Spirulina for the low-fat diet was obtained (Amazon.com, B0039ITKS8) and kept at room temperature until feeding. The high-fat diet was prepared according to (87). To obtain 8 % cholesterol supplementation, 80 mg of cholesterol was dissolved in 100 % ethanol under constant shaking. 1 g GEMMA Micro 150 (Skretting USA) was added and mixed until dissolved. After the evaporation of the ethanol overnight, the diet was ground through a 150-micron sieve to ensure the correct size of food for the consumption of the juvenile zebrafish. The diet was stored at 4 °C during the experiment.

### Feeding paradigm

After 5 dpf, 80 *apoBb.1^Dendra2/Dendra2^* were moved into a 10 L tank and 20 *apoBb.1^+/+^* larvae were moved into a 3 L tank. For apoC2 mutant feeding experiments, *apoC2^?/?^; apoBb.1^Dendra2/ Dendra2^* were housed in 10 L tanks together with *apoC2^?/?^; apoBb.1^+/+^* from 5 dpf until 14 dpf. All tanks were fed GEMMA Micro 75 (Skretting USA) three times a day until 14 dpf. At 14 dpf, larvae received 3 feedings of Western diet (Sparos: (Fish protein hydrolyzate 10 %; Cod powder 30 %; Fish gelatin 5 %; Pea Protein 7.5 %; Wheat gluten 7.5 %, Soybean Oil 7.5 %; Palm oil 8 %; Soy lecithin 4 %; Vit & Min Premix 2 %; Vitamin C 0.1 %; Cholesterol 4 %; Antioxidant 0.4 %; Monocalcium phosphate 3 %; Calcium silicate 2.2 %; L-Taurine 0.8 %; Fructose 8 %)). After a morning feeding with the Western diet, the juveniles were removed, and 36 *apoBb.1^Dendra2/ Dendra2^,* as well as 12 *apoBb.1^+/+^* animals, were selected. Animals were photo-converted in four batches of 9 *apoBb.1^Dendra2/Dendra2^* and 3 *apoBb.1^+/+^* animals. The juveniles were transferred into 48 playpens (92) in a total of four 10 L tanks, each 10 L tank housing 12 playpens. Going forward, two 10 L playpen-tanks were fed the low-fat diet, and the other two 10 L playpen-tanks received the high-fat diet. Every 24 hours, juveniles were removed 1.5 hours after the morning feed for imaging. The playpens were cleaned before the juveniles were returned after imaging. Juveniles were fed the second and third feeding of the day at normal times. Images were acquired until no more red-state ApoB-Dendra2 could be detected, and then juveniles were sacrificed. The playpens and false bottom were rinsed, cleaned in a Virkon (Syndel, VIRK-D-LB-0010) bath, and air-dried before being used again.

### Image processing for figures

All fluorescent images were adjusted for the intensity of the displayed channel using FIJI. Green channel images were adjusted to 50 - 1500. Images taken in the red channel were adjusted to 50-500 in Figure 2 and 50 - 300 in Figure 3 onwards. UV-channel images were set to 50 - 2000. To align larvae, images were rotated and cropped in Adobe Illustrator. The resulting data-free white space was filled in with black boxes when needed for visual ease. Images of the Western blot images were set to 0 - 0.1 and 0.002 intensity to allow visualization of all bands.

### Statistics

GraphPad Prism (GraphPad Software) was used for graphing and statistical analysis. Outliers in all datasets were identified by ROUT and excluded from the analysis. Brown-Forsythe and Welch ANOVA tests with Dunnett’s T3 multiple comparisons test were performed on half-life measures. Figure legends include sample sizes and details of statistical analysis.

---

**Figure S1:**
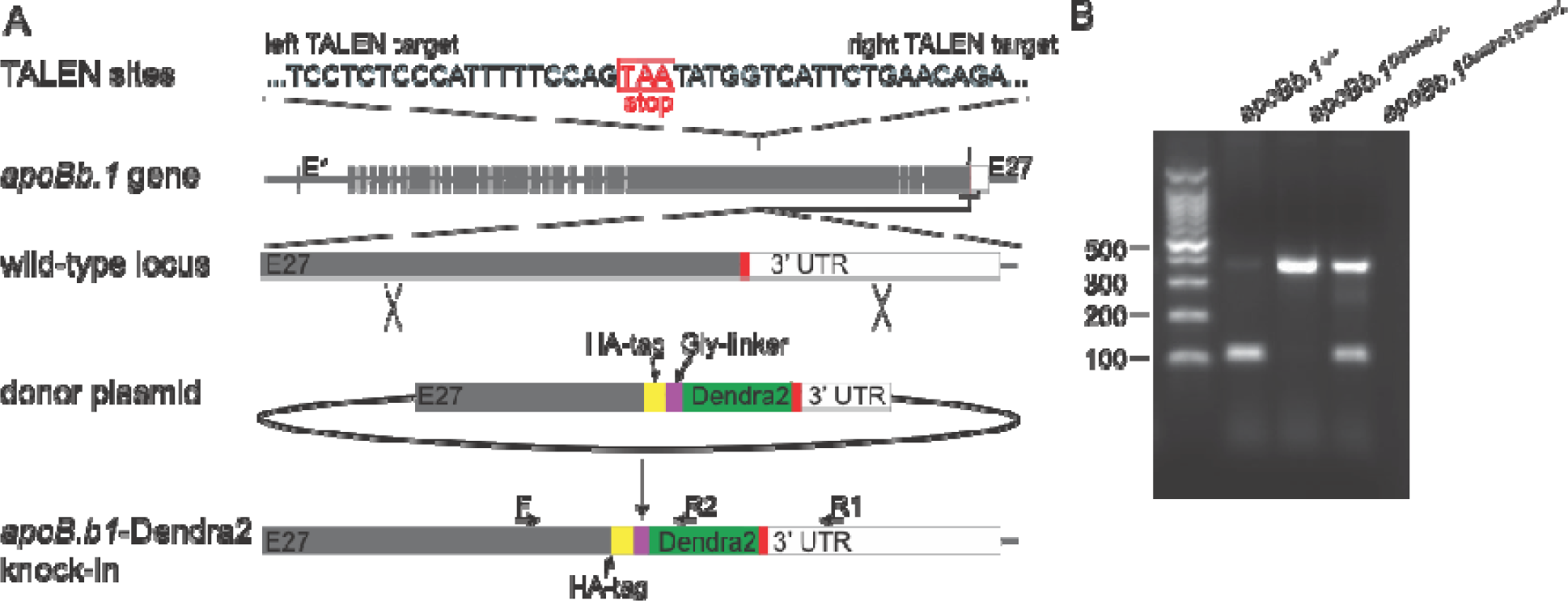
Generation of the in-frame fusion of Dendra2 to apoBb.1 using TALENs. (A) TALEN arms were previously published and evaluated for integration efficiency. The donor plasmid included TALEN cutting site and the sequences for an HA-tag followed by Dendra2. (B) PCR using primers 2-4 (S Table 4) which are located as indicated in (A) shows successful integration of Dendra2.

**Figure S2:**
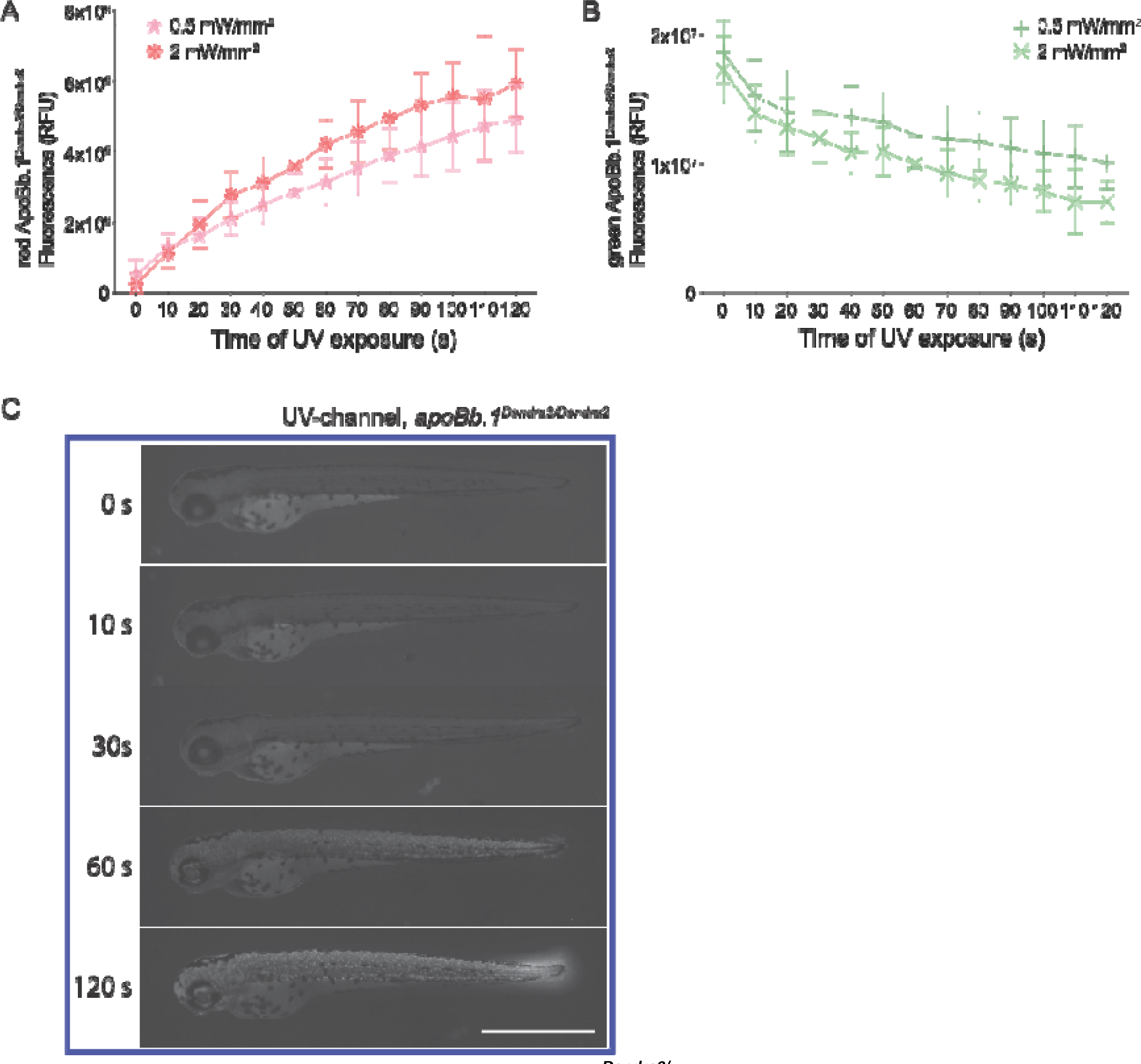
Incremental photoconversion in *apoBb.1^Dendra2/+^*larva and demonstration of tissue damage caused by exposure to UV light. (A+B) Three dpf old *apoBb.1^Dendra2/+^*larva were subjected to UV light of 0.5 or 2 mW/mm^2^ intensity in 10 s increments until 120 s. (A) Longer exposure and higher intensity increased the levels of red-state ApoB-Dendra2, while simultaneously decreasing (B) green-state ApoB-Dendra2. (C) In addition to imaging the green- and red-channel every 10 s, images were taken in the UV channel (excitation 395nm), revealing severe tissue abnormalities starting at 60 s of UV light exposure, which intensified after 120 s total exposure of UV light. Three independent experiments, n = 15. Scale bar 1 mm.

**Figure S3:**
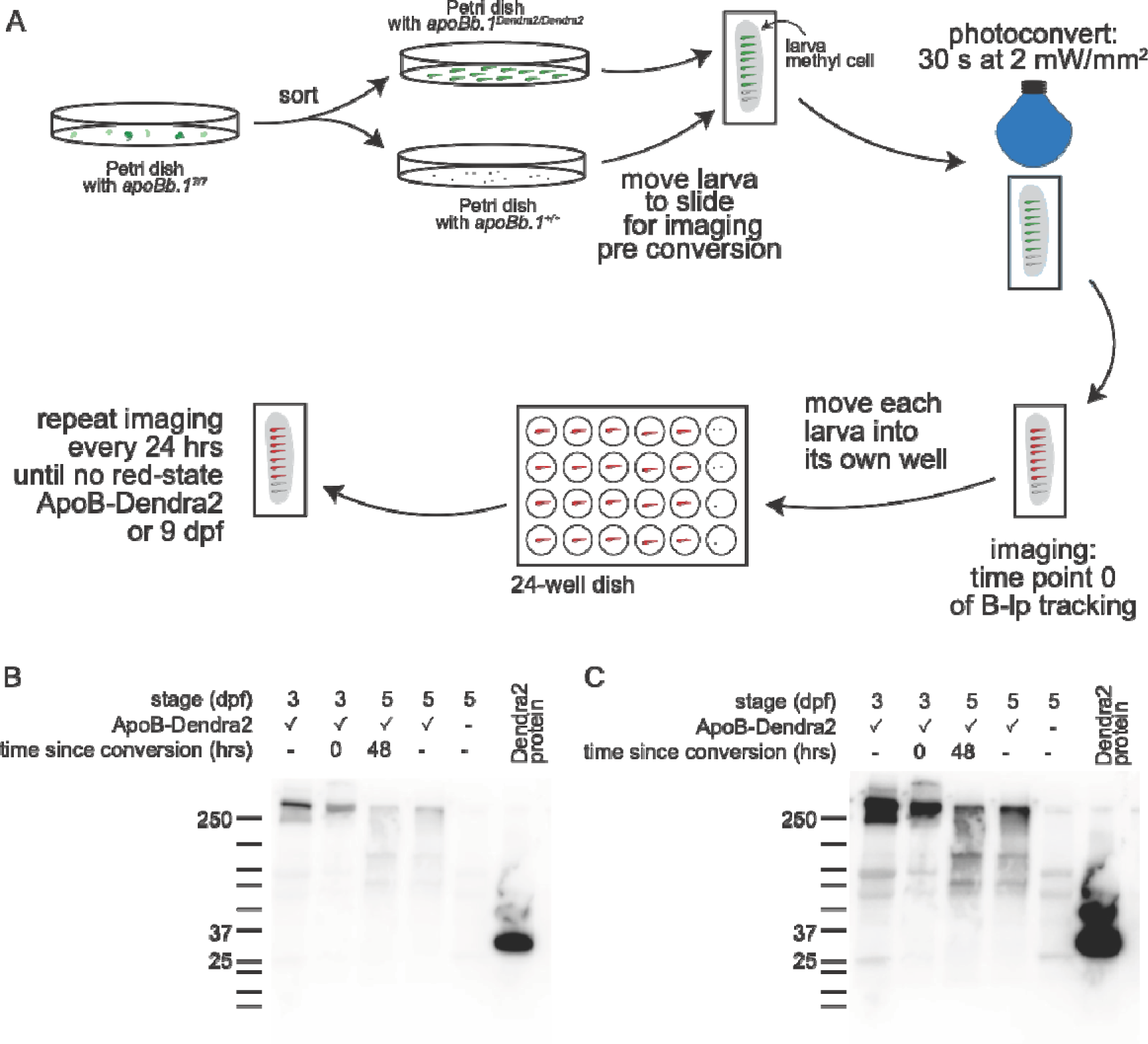
Experimental set up to measure B-lp half-life and Western blot confirming ApoB-Dendra2 stability. (A) Larva were sorted for *apoBb.1^Dendra2/Dendra2^* and *apoBb.1^+/+^* at 1 dpf, then raised in separate Petri dishes until the day of photoconversion. On the day of photoconversion, larvae were imaged, pre and post-photoconversion, then transferred into a well on a 24-well dish. Images were taken every 24 hrs of the same larvae. (B+C) Western blot shows that Dendra2 remains attached to ApoB, and its stability is not impacted by photoconversion. In each condition, 10 larvae (*apoBb.1^Dendra2/+^* or *apoBb1^+/+^*) were pooled for protein extraction. The blot was probed for Dendra2, and Dendra2-specific bands are only visible at 250 kDa, corresponding to the migration of ApoB. Dendra2 protein was purchased to show the migration pattern of free Dendra2 (about 27 kDa). FIJI exposure was set to show (B) Dendra2 and (C) all other bands. Three independent experiments.

**Figure S4:**
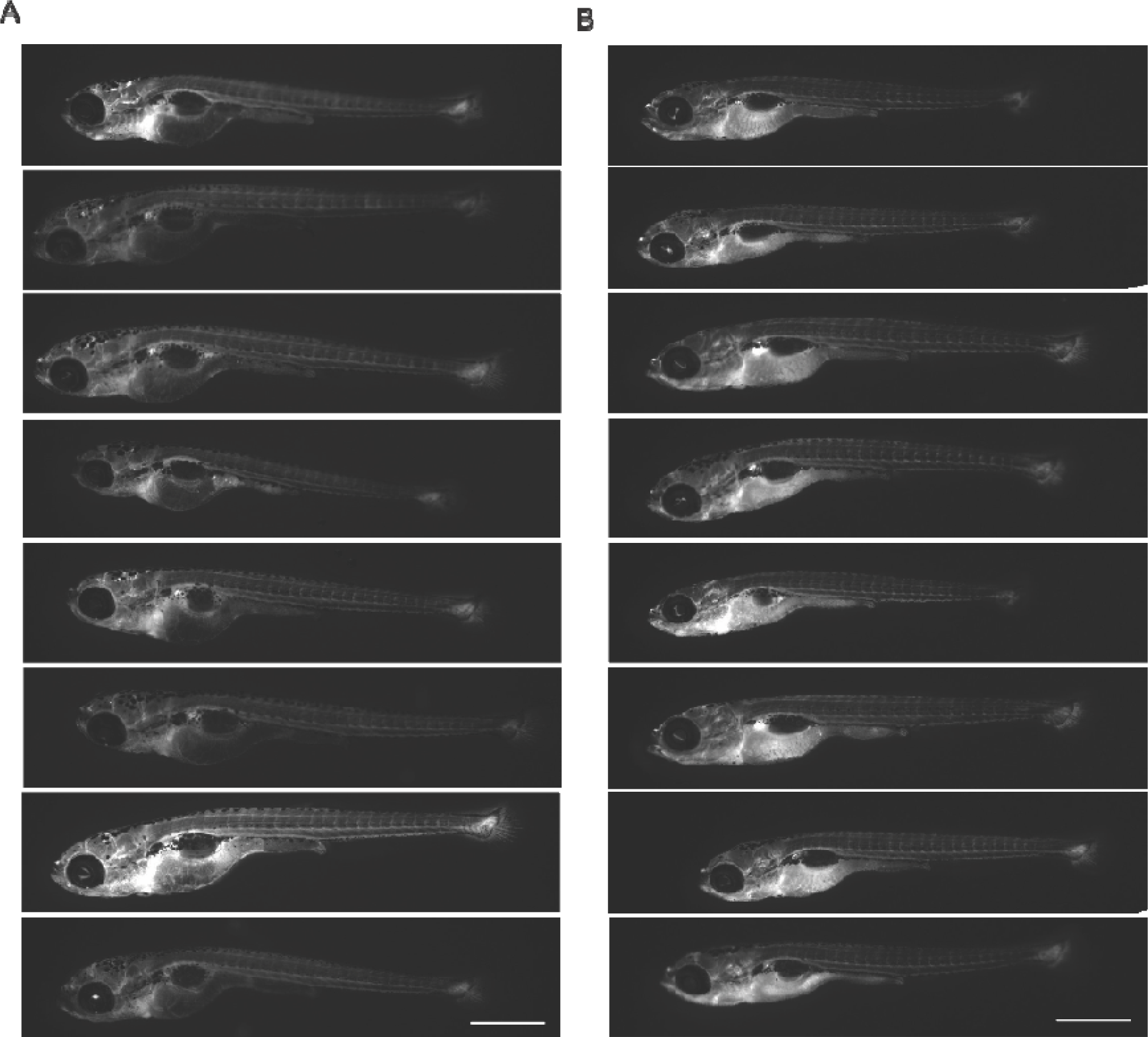
Feeding a Western diet for 1 day to achieve consistent B-lp levels for photoconversion. (A) Conventional feeding with GEMMA Micro 75 and GEMMA Micro 150 leads to varying levels of ApoB-Dendra2 B-lps in 15 dpf juveniles. (B) Feeding a Western diet for 1 day, results in homogenous increased levels of ApoB-Dendra2 B-lps. N = 36-40. Scale bar 1 mm.

**Figure S5:**
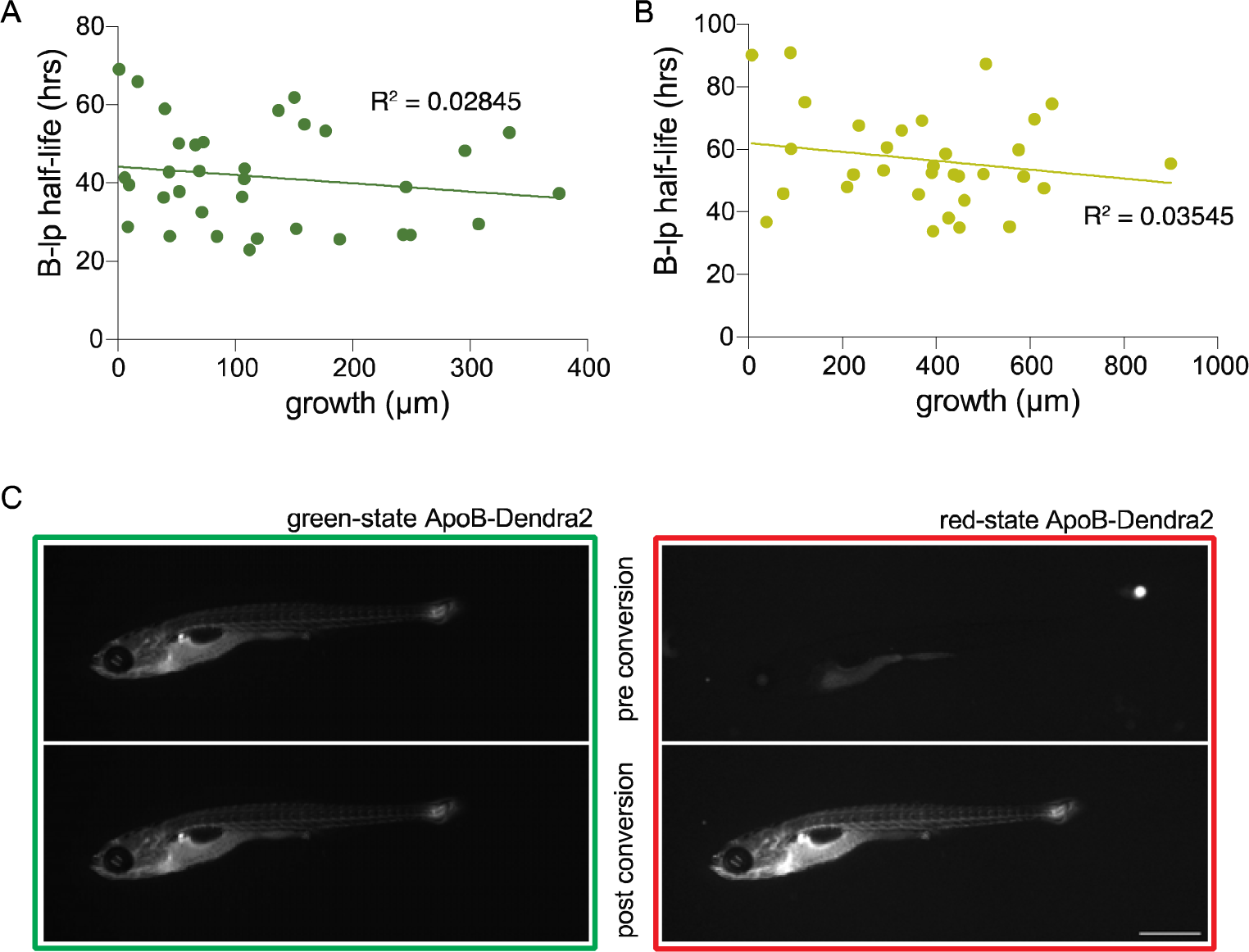
B-lp half-life does not correlate with the total growth of juveniles. Graphing B-lp turnover time in each animal in relation to its overall growth during the experiment. There is no correlation between the growth rate and the B-lp half-life of each individual larva when fed the (A) low fat diet and (B) high fat diet. The fitting of a trend line for the evaluation of correlation revealed a very low R^2^ in both diet conditions. Three independent experiments, n = 32-34. (C) Representative image of a juvenile wild-type zebrafish pre- and post-conversion highlighting the presence of chylomicrons in the intestinal folds that are readily photoconverted. Scale bar 1000 µm.

**Table S1:**
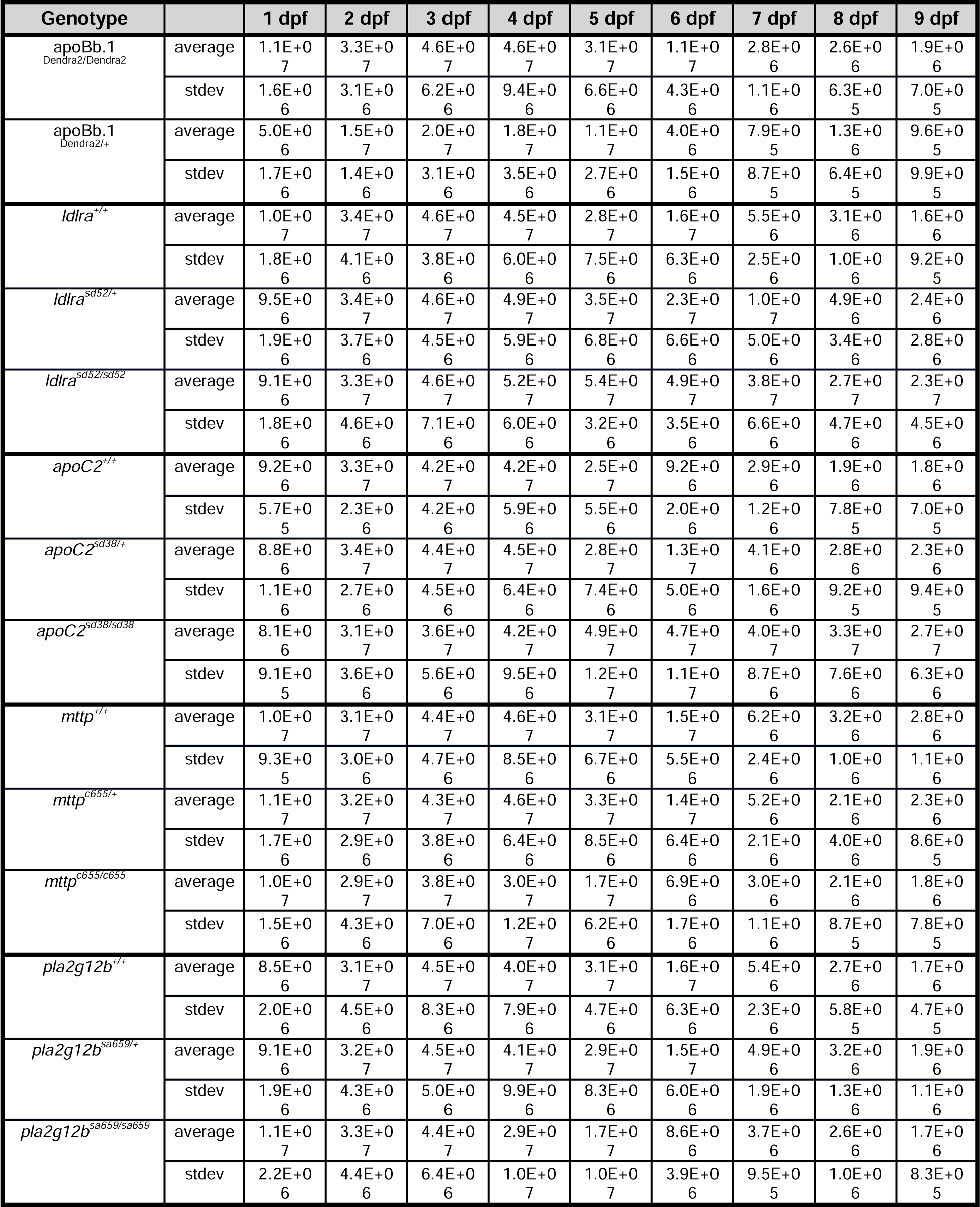
green-state ApoB-Dendra2 fluorescent values after background subtraction (in RFU, relative fluorescent units)

**Table S2:** Statistical analysis results of whole-body B-lp half-lives.

| Genotype | 2 dpf | 3 dpf | 4 dpf | 5 dpf |
| --- | --- | --- | --- | --- |
| *ldlra*<sup>+/+</sup> vs. *ldlra*<sup>sd52/+</sup> |  | 0.0002 | 0.2626 | 0.3654 |
| *ldlra*<sup>+/+</sup> vs. *ldlra*<sup>sd52/sd52</sup> |  | 0.0004 | <0.0001 | 0.0106 |
| *ldlra*<sup>sd52/+</sup> vs. *ldlra*<sup>sd52/sd52</sup> |  | 0.0890 | <0.0001 | 0.0251 |
| *apoC2*<sup>+/+</sup> vs. *apoC2*<sup>sd38/+</sup> |  | 0.1584 | 0.9895 | 0.5965 |
| *apoC2*<sup>+/+</sup> vs. *apoC2*<sup>sd38/sd38</sup> |  | <0.0001 | <0.0001 | 0.0011 |
| *apoC2*<sup>sd38/+</sup> vs. *apoC2*<sup>sd38/sd38</sup> |  | <0.0001 | <0.0001 | 0.0014 |
| *mttp*<sup>+/+</sup> vs. *mttp*<sup>c655/+</sup> | 0.5626 | 0.2332 | 0.6802 |  |
| *mttp*<sup>+/+</sup> vs. *mttp*<sup>c655/c655</sup> | 0.0198 | <0.0001 | 0.0932 |  |
| *mttp*<sup>c655/+</sup> vs. *mttp*<sup>c655/c655</sup> | 0.0631 | <0.0001 | 0.0617 |  |
| *pla2g12b*<sup>+/+</sup> vs. *pla2g12b*<sup>sa659/+</sup> | 0.9884 | 0.3694 | 0.6535 |  |
| *pla2g12b*<sup>+/+</sup> vs. *pla2g12b*<sup>sa659/sa659</sup> | 0.2050 | 0.0017 | 0.0357 |  |
| *pla2g12b*<sup>sa659/+</sup> vs. *pla2g12b*<sup>sa659/sa659</sup> | 0.1269 | 0.0006 | 0.0012 |  |

**Table S3:** Statistical analysis results of circulating B-lp half-lives.

| Genotype | 2 dpf | 3 dpf | 4 dpf |
| --- | --- | --- | --- |
| *mttp*<sup>+/+</sup> vs. *mttp*<sup>c655/+</sup> | 0.5610 | 0.8405 | 0.7078 |
| *mttp*<sup>+/+</sup> vs. *mttp*<sup>c655/c655</sup> | 0.0186 | 0.0028 | 0.0232 |
| *mttp*<sup>c655/+</sup> vs. *mttp*<sup>c655/c655</sup> | 0.0483 | 0.0001 | 0.0516 |
| *pla2g12b*<sup>+/+</sup> vs. *pla2g12b*<sup>sa659/+</sup> | 0.9421 | 0.4607 | 0.5981 |
| *pla2g12b*<sup>+/+</sup> vs. *pla2g12b*<sup>sa659/sa659</sup> | 0.0798 | 0.0020 | 0.0202 |
| *pla2g12b*<sup>sa659/+</sup> vs. *pla2g12b*<sup>sa659/sa659</sup> | 0.0562 | 0.0027 | 0.0048 |

**Table S4:**
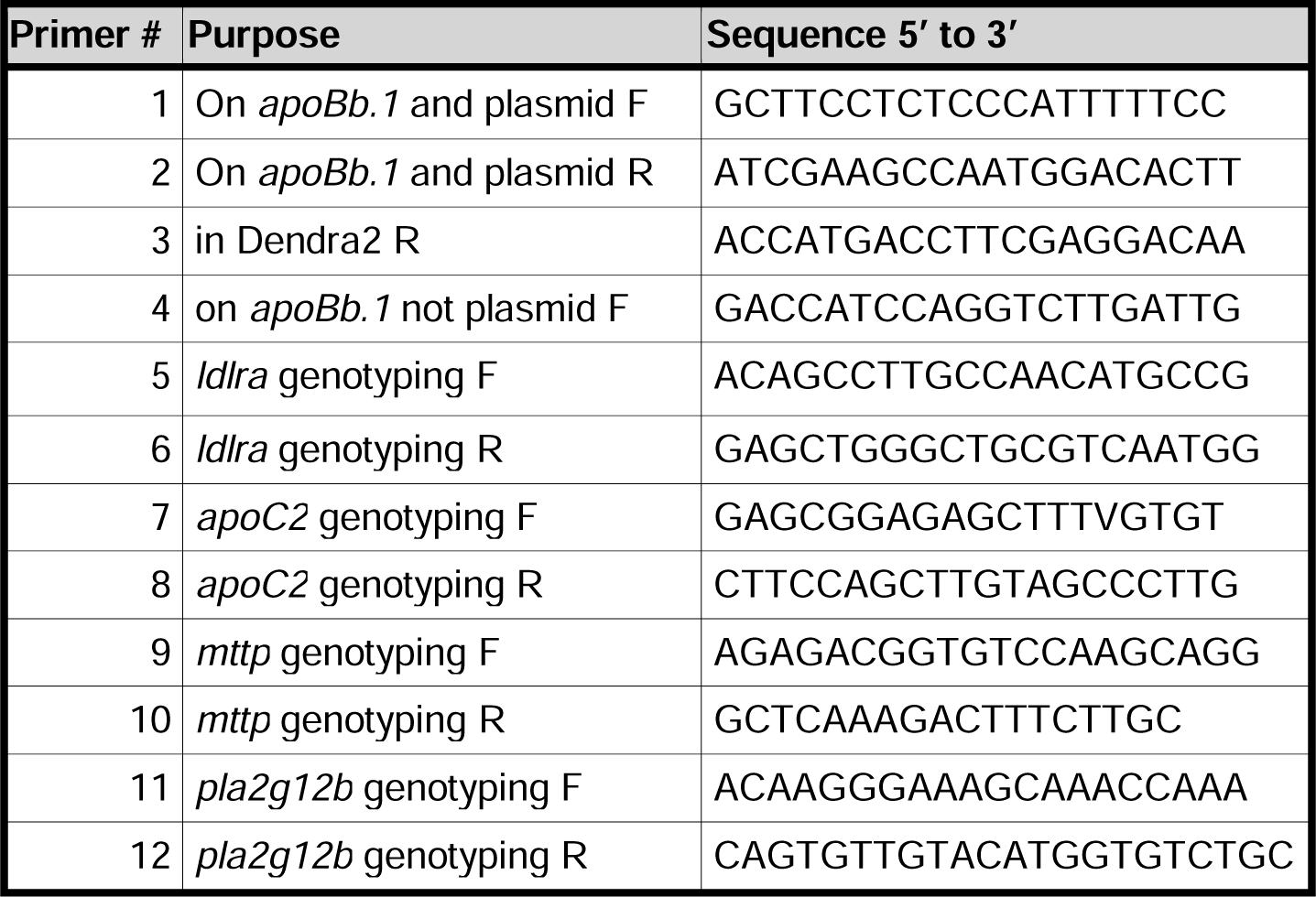
Primers used in this study.

